# Unconventional kinetochore kinases KKT2 and KKT3 have unique centromere localization domains

**DOI:** 10.1101/2019.12.13.875419

**Authors:** Gabriele Marcianò, Midori Ishii, Olga O. Nerusheva, Bungo Akiyoshi

**Author notes:** These authors contributed equally to this work (Gabriele Marcianò, Midori Ishii, Olga O. Nerusheva).

## Abstract

Chromosome segregation in eukaryotes is driven by the kinetochore, the macromolecular protein complex that assembles onto centromeric DNA and binds spindle microtubules. Cells must tightly control the number and position of kinetochores so that all chromosomes assemble a single kinetochore. A central player in this process is the centromere-specific histone H3 variant CENP-A, which localizes constitutively at centromeres and promotes kinetochore assembly. However, CENP-A is absent from several eukaryotic lineages including kinetoplastids, a group of evolutionarily divergent eukaryotes that have an unconventional set of kinetochore proteins. There are six proteins that localize constitutively at centromeres in the kinetoplastid parasite *Trypanosoma brucei,* among which two homologous protein kinases (KKT2 and KKT3) have limited similarity to polo-like kinases. In addition to the N-terminal kinase domain and the C-terminal divergent polo boxes, KKT2 and KKT3 have a central domain of unknown function as well as putative DNA-binding motifs. Here we show that KKT2 and KKT3 are important for the localization of several kinetochore proteins and that their central domains are sufficient for centromere localization in *T. brucei*. Crystal structures of the KKT2 central domain from two divergent kinetoplastids reveal a unique zinc-binding domain (termed the CL domain for centromere localization), which promotes its kinetochore localization in *T. brucei*. Mutations in the equivalent domain in KKT3 abolish its kinetochore localization and function. Our work shows that the unique central domains play a critical role in mediating the centromere localization of KKT2 and KKT3.

## Introduction

The kinetochore is the macromolecular protein complex that drives chromosome segregation during mitosis and meiosis in eukaryotes. Its fundamental functions are to bind DNA and spindle microtubules (Musacchio and Desai, 2017). In most eukaryotes, kinetochores assemble within a single chromosomal region called the centromere. While components of spindle microtubules are highly conserved across eukaryotes (Wickstead and Gull, 2011; Findeisen et al., 2014), centromere DNA is known to evolve rapidly (Henikoff et al., 2001). Nevertheless, it is critical that a single kinetochore is assembled per chromosome and its position is maintained between successive cell divisions. A key player involved in this kinetochore specification process is the centromere-specific histone H3 variant CENP-A, which is found in most sequenced eukaryotic genomes (Talbert et al., 2009). CENP-A localizes specifically at centromeres throughout the cell cycle and recruits HJURP, a specific chaperone that incorporates CENP-A onto centromeres (Black and Cleveland, 2011; McKinley and Cheeseman, 2016; Stankovic and Jansen, 2017). Besides CENP-A, components of the constitutive centromere-associated network (CCAN) also localize at centromeres throughout the cell cycle. CENP-A-containing nucleosomes are recognized by CCAN components, which in turn recruit the KNL1-Mis12-Ndc80 (KMN) network that has microtubule-binding activities. In addition to these structural kinetochore proteins, several protein kinases are known to localize at mitotic kinetochores, including Cdk1, Aurora B, Bub1, Mps1, and Plk1 (Cheeseman and Desai, 2008). These protein kinases regulate various aspects of mitosis, including kinetochore assembly, error correction, and the spindle checkpoint (Carmena et al., 2012; London and Biggins, 2014; Hara and Fukagawa, 2018).

Kinetoplastids are evolutionarily divergent eukaryotes that are defined by the presence of a unique organelle called the kinetoplast that contains a cluster of mitochondrial DNA (d’Avila-Levy et al., 2015). Centromere positions have been mapped in three kinetoplastids: 20–120 kb regions that have AT-rich repetitive sequences in *Trypanosoma brucei* (Obado et al., 2007), ~16 kb GC-rich unique sequences in *Trypanosoma cruzi* (Obado et al., 2005), and ~4 kb regions in *Leishmania major* (Garcia-Silva et al., 2017). Although some DNA elements and motifs are enriched, there is no specific DNA sequence that is common to all centromeres in each organism, suggesting that kinetoplastids likely determine their kinetochore positions in a largely sequence-independent manner. However, none of CENP-A or any other canonical structural kinetochore protein has been identified in kinetoplastids (Lowell and Cross, 2004; Berriman et al., 2005; Aslett et al., 2010). They instead have unique kinetochore proteins, such as KKT1–25 (Akiyoshi and Gull, 2014; Nerusheva and Akiyoshi, 2016; Nerusheva et al., 2019) and KKIP1–12 (D’Archivio and Wickstead, 2017; Brusini et al., 2019) in *T. brucei*. It remains unknown which of these proteins form the base of trypanosome kinetochores that recruits other proteins. There are six proteins that localize at centromeres throughout the cell cycle (KKT2, KKT3, KKT4, KKT20, KKT22, and KKT23), implying their close association with centromeric DNA. Indeed, we previously showed that KKT4 has DNA-binding activity in addition to microtubule-binding activity (Llauró et al., 2018; Ludzia et al., 2020). However, RNAi-mediated knockdown of KKT4 affected the localization of KKT20 but not other kinetochore proteins, suggesting that KKT4 is largely dispensable for kinetochore assembly.

In this study, we focused on KKT2 and KKT3, which are homologous to each other and have three domains conserved among kinetoplastids: a protein kinase domain that is classified as unique among known eukaryotic kinase subfamilies (Parsons et al., 2005), a central domain of unknown function, and divergent polo boxes (DPB). Presence of an N-terminal kinase domain and a C-terminal DPB suggests that KKT2 and KKT3 likely share common ancestry with polo-like kinases (Nerusheva and Akiyoshi, 2016). Interestingly, a protein kinase domain is not present in any constitutively-localized kinetochore protein in other eukaryotes, making these protein kinases a unique feature of kinetoplastid kinetochores. In addition to the three domains that are highly conserved among kinetoplastids, AT-hook and SPKK DNA-binding motifs are found in some species, suggesting that these proteins are located close to DNA (Akiyoshi and Gull, 2014). Although RNAi-mediated knockdown of KKT2 or KKT3 leads to growth defects (Akiyoshi and Gull, 2014; Jones et al., 2014), little is known about their molecular function. In this report, we have revealed a unique zinc-binding domain in the KKT2 and KKT3 central domain, which is important for their kinetochore localization and function in *T. brucei*.

## Results

### Localization of KKT2 and KKT3 is not affected by depletion of various kinetochore proteins

We previously showed in *T. brucei* procyclic form (insect stage) cells that kinetochore localization of KKT2 and KKT3 was not affected by KKT4 depletion (Llauró et al., 2018). To examine the effect of other kinetochore proteins for the recruitment of KKT2 and KKT3, we established RNAi-mediated knockdowns for KKT1, KKT6, KKT7, KKT8, KKT10/19, KKT14, KKT22, KKT23, KKT24, and KKIP1 (Figure 1A). Using inducible stem-loop RNAi constructs in cells expressing YFP-fusion of the target protein, we confirmed efficient depletion of YFP signals and observed severe growth defects for KKT1, KKT6, KKT8, KKT14, and KKT24. RNAi against KKT7, KKT10/19, and KKIP1 also caused severe growth defects as previously observed (D’Archivio and Wickstead, 2017; Ishii and Akiyoshi, 2020). In contrast, only a mild growth defect was observed for KKT23, while no growth defect was observed for KKT22. We next used these RNAi constructs in cells expressing either YFP-KKT2 or KKT3-YFP and found that these proteins formed kinetochore-like dots in all conditions (Figure 1B, C). These results show that KKT2 and KKT3 can localize at kinetochores even when various kinetochores proteins are depleted.

**Figure 1.**
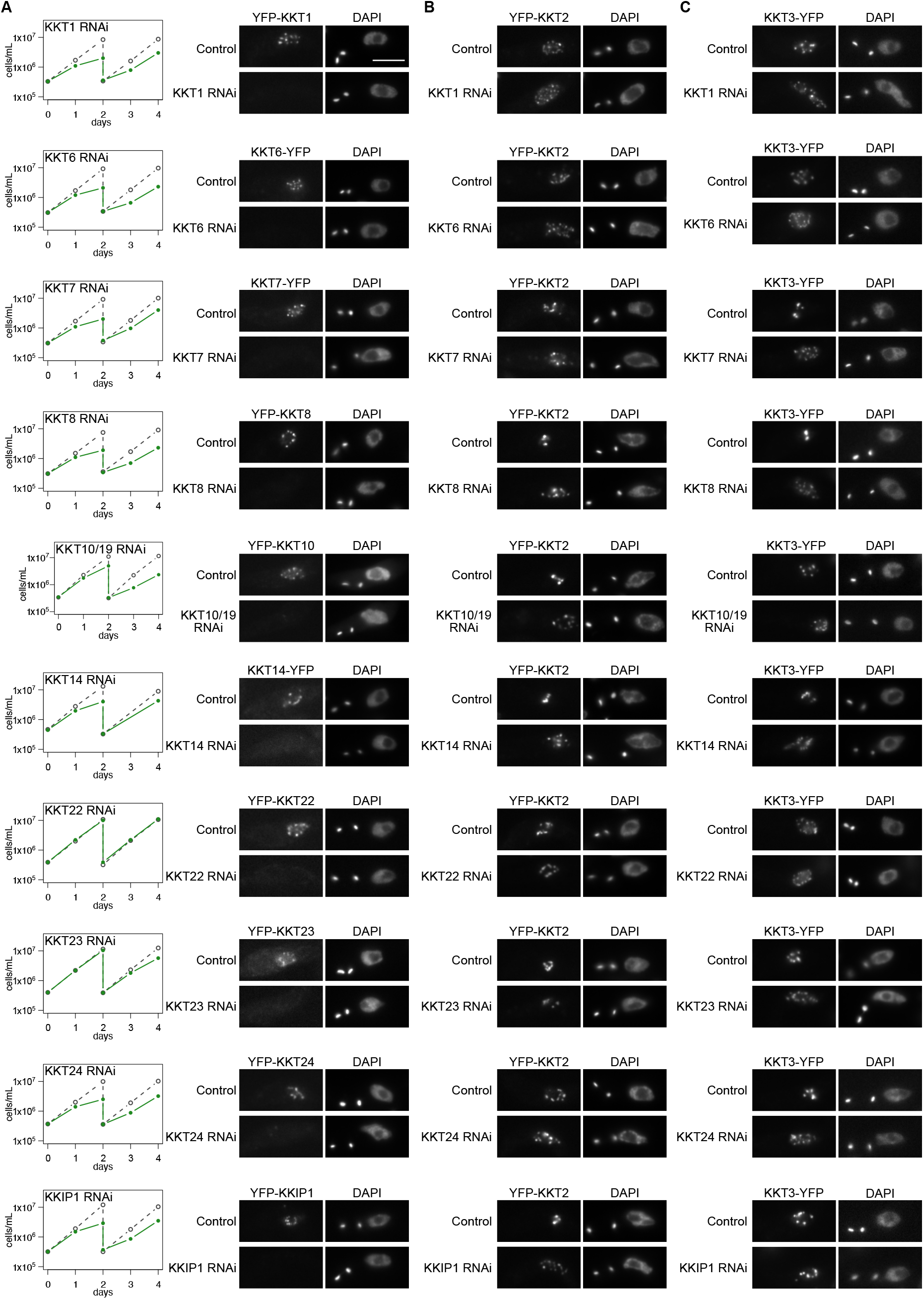
Kinetochore localization of KKT2 and KKT3 is not dependent on various kinetochore proteins. (A) Left: Growth curves of control (gray dashed line) and RNAi-induced cultures (green line) show that these kinetochore proteins are important for proper cell growth (except KKT22) in *T. brucei* procyclic form cells. RNAi was induced with 1 μg/mL doxycycline. Cultures were diluted at day 2. Right: Depletion of indicated kinetochore proteins was confirmed by microscopy. Cells were fixed at either day 1 (RNAi against KKT1, KKT6, KKT7 5′UTR, KKT8 3′UTR, KKT10/19, KKT14, KKT24, and KKIP1) or day 2 (RNAi against KKT22 and KKT23). *T. brucei* has a kinetoplast (K) that contains mitochondrial DNA, and a nucleus (N) that contains nuclear DNA. Kinetoplasts segregate prior to the nuclear division, and the number of kinetoplasts and nuclei can be used as a cell cycle marker (Woodward and Gull, 1990; Siegel et al., 2008). YFP signal was efficiently depleted in > 60% of 2K1N cells (G2 to metaphase) in each case (n > 50, each). Cell lines, BAP672, BAP699, BAP2001, BAP2002, BAP139, BAP2086, BAP1840, BAP1842, BAP1843, BAP770. (B) Kinetochore localization of KKT2 is not affected by depletion of various kinetochore proteins. RNAi against indicated kinetochore proteins was induced in cells expressing YFP-KKT2. Dot formation was observed in >94% of 2K1N cells in all cases (n > 70, each). Cells were fixed as in (A), except for KKT23 RNAi cells that were fixed at day 4. Cell lines, BAP2004, BAP2005, BAP2006, BAP2007, BAP2008, BAP680, BAP2010, BAP2011, BAP2012, BAP2013. (C) Kinetochore localization of KKT3 is not affected by depletion of various kinetochore proteins. RNAi against indicated kinetochore proteins was induced in cells expressing KKT3-YFP. Dot formation was observed in >97% of 2K1N cells in all cases (n > 50, each). Cells were fixed as in (B). Cell lines, BAP2014, BAP2015, BAP2016, BAP2017, BAP2018, BAP2085, BAP2020, BAP2021, BAP2022, BAP2023. Scale bars, 5 μm.

### KKT2 and KKT3 are important for localization of some kinetochore proteins

We next examined whether KKT2 and KKT3 are important for the localization of other kinetochore proteins. KKT2 RNAi using a stem-loop construct caused growth defects, as previously reported (Figure 2A) (Ishii and Akiyoshi, 2020). We saw defective kinetochore localization for KKT14 upon induction of KKT2 RNAi, while other tested proteins still formed kinetochore-like dots at 1 day post-induction (Figure 2B, C). We next established RNAi against KKT3 (Figure 2D) and examined its effect on the localization of other kinetochore proteins at 2 days post-induction. Kinetochore localization of two constitutive kinetochore components, KKT22 and KKT23, was affected by KKT3 depletion (Figure 2E, F), while that of other tested proteins remained largely intact. It is noteworthy that KKT2 and KKT3 can apparently localize at kinetochores independently from each other. Together with the fact that KKT2 and KKT3 are homologous proteins, these results raised a possibility that KKT2/3 might have redundant roles in kinetochore assembly, so we next examined the effect of double knockdown. Although the growth defect of the KKT2/3 double RNAi was not dramatically different from that of individual knockdowns (Figure 2G), which could be explained by residual KKT2/3 signals (Figure 2I), defective kinetochore localization was found for KKT1 and KKT4 at 1 day post-induction (Figure 2H, I). Taken together, these results show that KKT2 and KKT3 play important roles in recruiting multiple kinetochore proteins. It is possible that more efficient or rapid inactivation methods could reveal additional kinetochore proteins whose localizations depend on KKT2/3.

**Figure 2.**
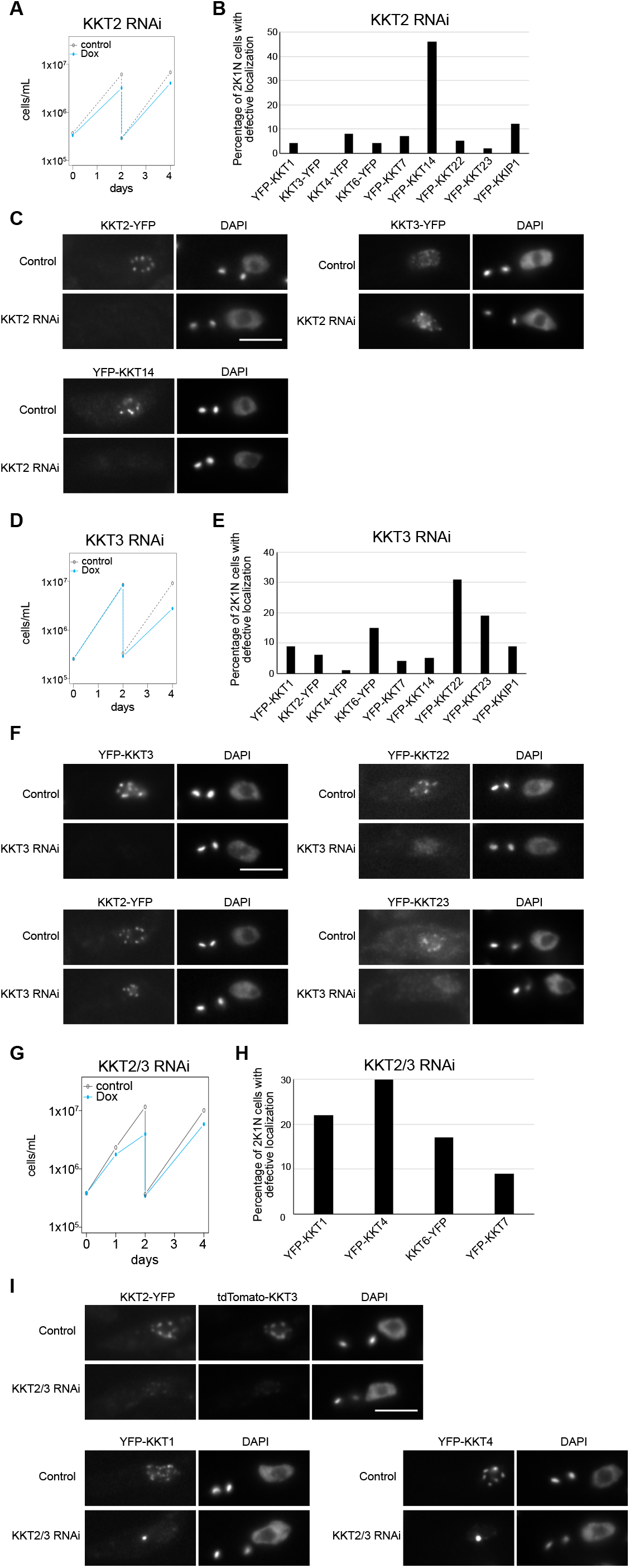
KKT2 and KKT3 are important for kinetochore assembly. (A) Growth curve for KKT2 5′UTR RNAi in cells expressing KKT2-YFP. RNAi was induced with 1μg/mL doxycycline and cultures were diluted at day 2. KKT2-YFP signal was depleted in 46% of 2K1N cells (G2 to metaphase) at day 1 post-induction (n > 100). Cell line, BAP1752. (B) Quantification of 2K1N cells that had defective kinetochore localization of indicated kinetochore proteins upon induction of KKT2 5′UTR RNAi (n > 100, each). Cells were fixed at day 1 post-induction. In each case, normal kinetochore signal was confirmed in un-induced 2K1N cells (not shown). Cell lines, BAP1743, BAP1753, BAP1749, BAP2075, BAP1750, BAP1746, BAP1751, BAP2080, BAP1747. (C) Examples of cells expressing indicated kinetochore proteins fused with YFP, showing that KKT3 still forms kinetochore-like dots, while KKT14 fails to localize at kinetochores upon KKT2 depletion. (D) Growth curve for KKT3 3′UTR RNAi in cells expressing YFP-KKT3. Cultures were diluted at day 2. YFP-KKT3 signal was depleted in 61% of 2K1N cells at day 2 post-induction (n > 100). Cell line, BAP1659. (E) Quantification of 2K1N cells that had defective kinetochore localization of indicated kinetochore proteins upon induction of KKT3 3′UTR RNAi for 2 days (n > 100, each). In each case, normal kinetochore signal was confirmed in un-induced 2K1N cells (not shown). Cell lines, BAP1755, BAP1764, BAP1761, BAP2076, BAP1762, BAP1758, BAP1763, BAP2081, BAP1759. (F) Examples of cells expressing indicated kinetochore proteins fused with YFP, showing that KKT2 still forms kinetochore-like dots, while KKT22 and KKT23 failed to localize at kinetochores upon KKT3 depletion. (G) Growth curve for KKT2/3 double RNAi that targets KKT2 5′UTR and KKT3 3′UTR in cells expressing KKT2-YFP and tdTomato-KKT3. Both KKT2 and KKT3 signals were depleted in 55% of 2K1N cells at day 1 post-induction (n > 100). Cell line, BAP2089. (H) Quantification of 2K1N cells that had defective kinetochore localization of indicated kinetochore proteins upon induction of KKT2/3 double RNAi for 1 day (n > 100, each). In each case, normal kinetochore signal was confirmed in un-induced 2K1N cells (not shown). Cell lines, BAP2068, BAP2070, BAP2074, BAP2071. (I) Examples of cells expressing indicated kinetochore proteins fused with fluorescent proteins, showing that KKT1 and KKT4 failed to form normal kinetochore-like dots but instead formed bright blobs upon KKT2/3 depletion. Cells were fixed at day 1 post-induction. Scale bars, 5 μm.

### Multiple domains of KKT2 are able to localize at centromeres in *T. brucei*

To understand how KKT2 localizes at centromeres, we determined which domain was responsible for its centromere localization by expressing a series of truncated versions of KKT2, fused with a GFP-tagged nuclear localization signal peptide (GFP-NLS) in *T. brucei* (Figure 3A). We previously showed that ectopically-expressed KKT2 DPB (residues 1024–1260) localized at kinetochores (Nerusheva and Akiyoshi, 2016). The present study confirmed this result and also identified two other regions (residues 562–677 and 672–1030) that localized at kinetochores from S phase to anaphase (Figure 3A, B and Figure S1).

**Figure 3.**
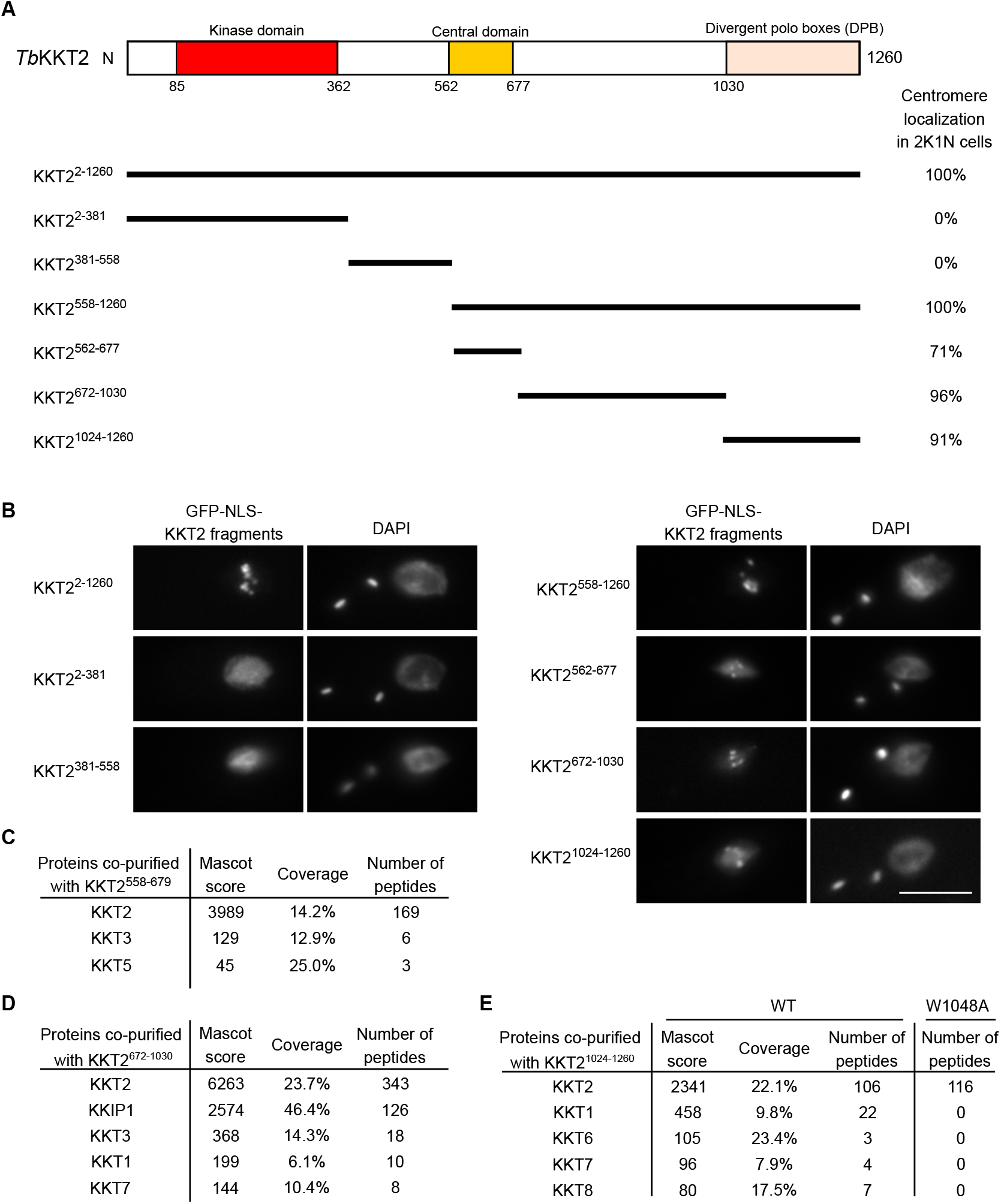
KKT2 has multiple domains that can promote centromere localization in *T. brucei*. (A) Schematic of the *T. brucei* KKT2 protein. Percentages of GFP-positive 2K1N cells (G2 to metaphase) that have kinetochore-like dots were quantified at 1 day post-induction (n > 22, each). (B) Ectopically expressed *Tb*KKT2 fragments that contain either the central domain (562–677), 672– 1030, or the divergent polo boxes (1024–1260) form kinetochore-like dots. Inducible GFP-NLS fusion proteins were expressed with 10 ng/mL doxycycline. Cell lines, BAP327, BAP328, BAP381, BAP331, BAP457, BAP519, BAP517. Scale bar, 5 μm. (C) *Tb*KKT2^558–679^ does not co-purify robustly with other kinetochore proteins. Cell line, BAP382. (D) *Tb*KKT2^672–1030^ co-purifies with KKIP1 and several other kinetochore proteins. Cell line, BAP519. (E) *Tb*KKT2 DPB WT, not W1048A, co-purifies with several kinetochore proteins. Cell lines, BAP517, BAP535. Inducible GFP-NLS fusion proteins were expressed with 10 ng/mL doxycycline, and immunoprecipitation was performed using anti-GFP antibodies. See Table S1 for all proteins identified by mass spectrometry.

Based on our previous finding that KKT2 co-immunoprecipitated with a number of kinetochore proteins (Akiyoshi and Gull, 2014), we reasoned that these fragments might localize at kinetochores by interacting with other kinetochore proteins. To test this possibility, we immunoprecipitated KKT2 fragments and performed mass spectrometry to identify co-purifying proteins. Although the central domain co-purified only with limited amounts of KKT3 and KKT5 (Figure 3C and Table S1), KKT2^672–1030^ co-purified with several kinetochore proteins, with KKIP1 being the top hit (Figure 3D and Table S1). Similarly, KKT2 DPB co-purified with several kinetochore proteins, which was abolished in the W1048A mutant that did not localize at kinetochores (Figure 3E and Table S1) (Nerusheva and Akiyoshi, 2016). These results support the possibility that ectopically-expressed KKT2^672–1030^ and DPB are able to localize at kinetochores from S phase to anaphase by interacting with non-constitutive kinetochore proteins (e.g. KKT1, KKT6, KKT7, KKT8, KKIP1). A corollary is that, in wild-type cells, the constitutively-localized KKT2 protein recruits these transient kinetochore proteins onto kinetochores using KKT2^672–1030^ and DPB domains.

### The central domain of KKT3 is able to localize at centromeres constitutively

We next expressed KKT3 fragments in trypanosomes (Figure 4A, B). Similarly to KKT2, the N-terminal protein kinase domain of KKT3 did not localize at centromeres. KKT3 DPB had robust kinetochore localization only during anaphase (Figure 4C), which differs from KKT2 DPB that localized from S phase to anaphase. Immunoprecipitation of KKT3 DPB identified a number of co-purifying kinetochore proteins (Figure 4D), raising a possibility that KKT3, like KKT2, recruits other kinetochore proteins by its DPB.

**Figure 4.**
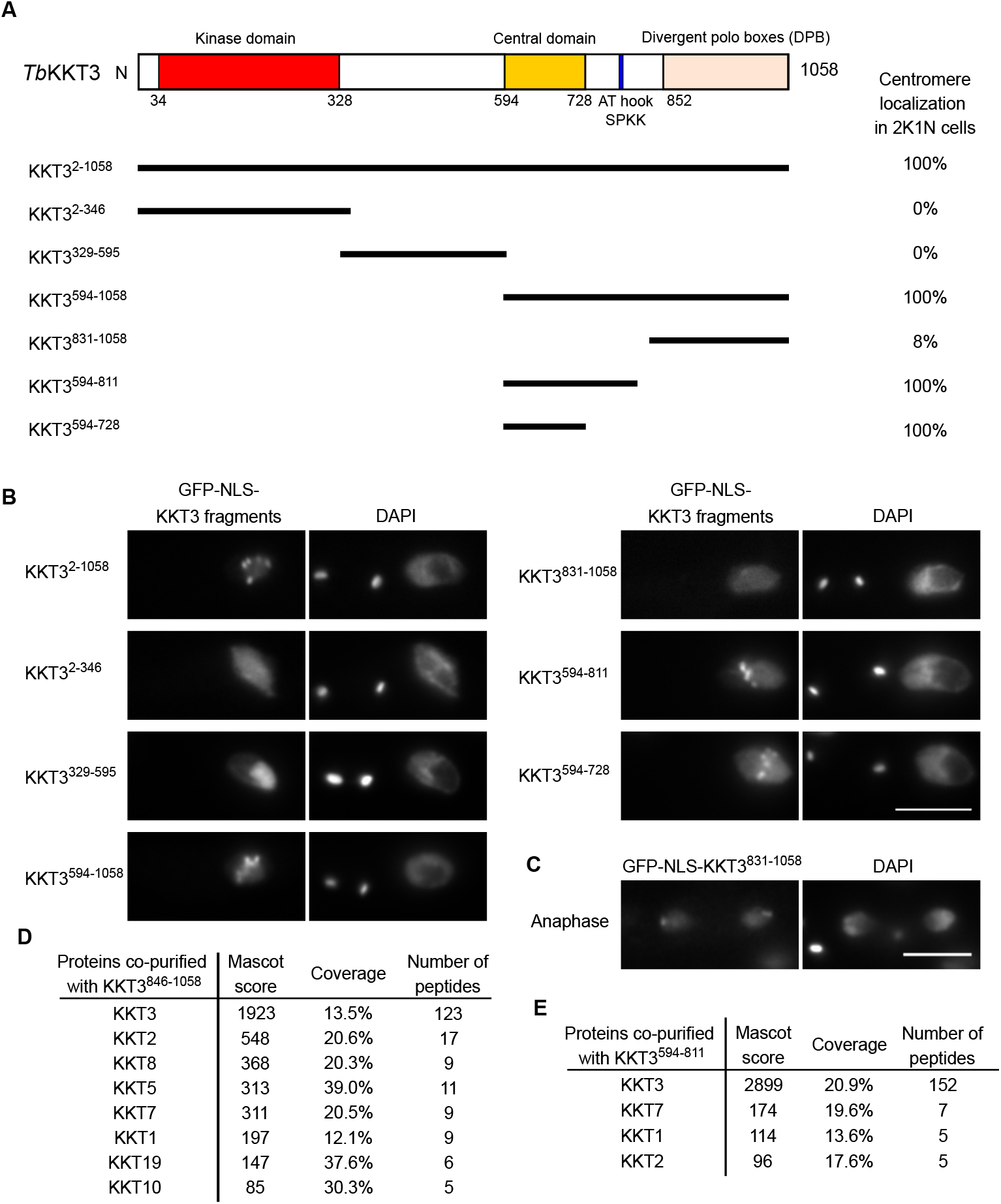
KKT3 central domain is able to localize at centromeres constitutively in *T. brucei*. (A) Schematic of the *T. brucei* KKT3 protein. Percentages of GFP-positive 2K1N cells (G2 to metaphase) that have kinetochore-like dots were quantified at 1 day post-induction (n > 24, each). (B) Ectopically expressed *Tb*KKT3 fragments that contain the central domain form kinetochore-like dots. Inducible GFP-NLS fusion proteins were expressed with 10 ng/mL doxycycline. Cell lines, BAP291, BAP292, BAP379, BAP296, BAP378, BAP377, BAP418. Scale bar, 5 μm. (C) *Tb*KKT3 DPB (831–1058) forms kinetochore-like dots during anaphase (88% of 2K2N cells, n = 25). Note that kinetochore proteins typically localize near the leading edge of separating chromosomes during anaphase. Cell line, BAP296. Scale bar, 5 μm. (D) *Tb*KKT3 DPB co-purifies with several kinetochore proteins. Cell line, BAP520. (E) *Tb*KKT3^594–811^ does not co-purify robustly with other kinetochore proteins. Cell line, BAP377. See Table S1 for all proteins identified by mass spectrometry.

KKT3^594–1058^ and KKT3^594–811^ that contain the central region also localized at kinetochores. Kinetochore localization was also observed for KKT3^594–728^, which lacks AT-hook and SPKK DNA-binding motifs (residues 771–780). KKT3^594–811^ co-purified with limited amounts of KKT7, KKT1, and KKT2 (Figure 4E). Importantly, KKT3^594–728^ constitutively localized at kinetochores (Figure S1), suggesting that the central domain plays a crucial role in recruiting KKT3 onto centromeres throughout the cell cycle.

### The *Bodo saltans* KKT2 central domain adopts a unique structure

To gain insights into how the central domains of KKT2 and KKT3 localize at centromeres, we expressed and purified recombinant proteins for their structure determination by X-ray crystallography. Our attempts to purify the *T. brucei* KKT3 (*Tb*KKT3 hereafter) central domain were unsuccessful, but we managed to express and purify from *E. coli* the central domain of KKT2 from several kinetoplastids, including *Bodo saltans* (a free-living kinetoplastid (Jackson et al., 2016)) and *Perkinsela* (endosymbiotic kinetoplastids (Tanifuji et al., 2017)) (Figure S2). We obtained crystals of *Bs*KKT2^572–668^ (which corresponds to residues 569–664 in *T. brucei* KKT2) and determined its structure to 1.8 Å resolution by zinc single-wavelength anomalous dispersion (Zn-SAD) phasing (Figure 5 and Table 1). Our analysis revealed the presence of two distinct zinc-binding domains: the N-terminal one (referred to as the “CL” domain for its key role in centromere localization: see below) consists of 2 β-sheets (where β-strands 1, 4 and 5 comprise the first β-sheet, and β-strands 2 and 3 comprise the second β-sheet) and one α-helix, while the C-terminal one consists of one β-sheet (comprising β-strands 6 and 7) and one α-helix (Figure 5A, B). The CL domain coordinates two zinc ions and the C-terminal domain coordinates one zinc ion.

**Figure 5.**
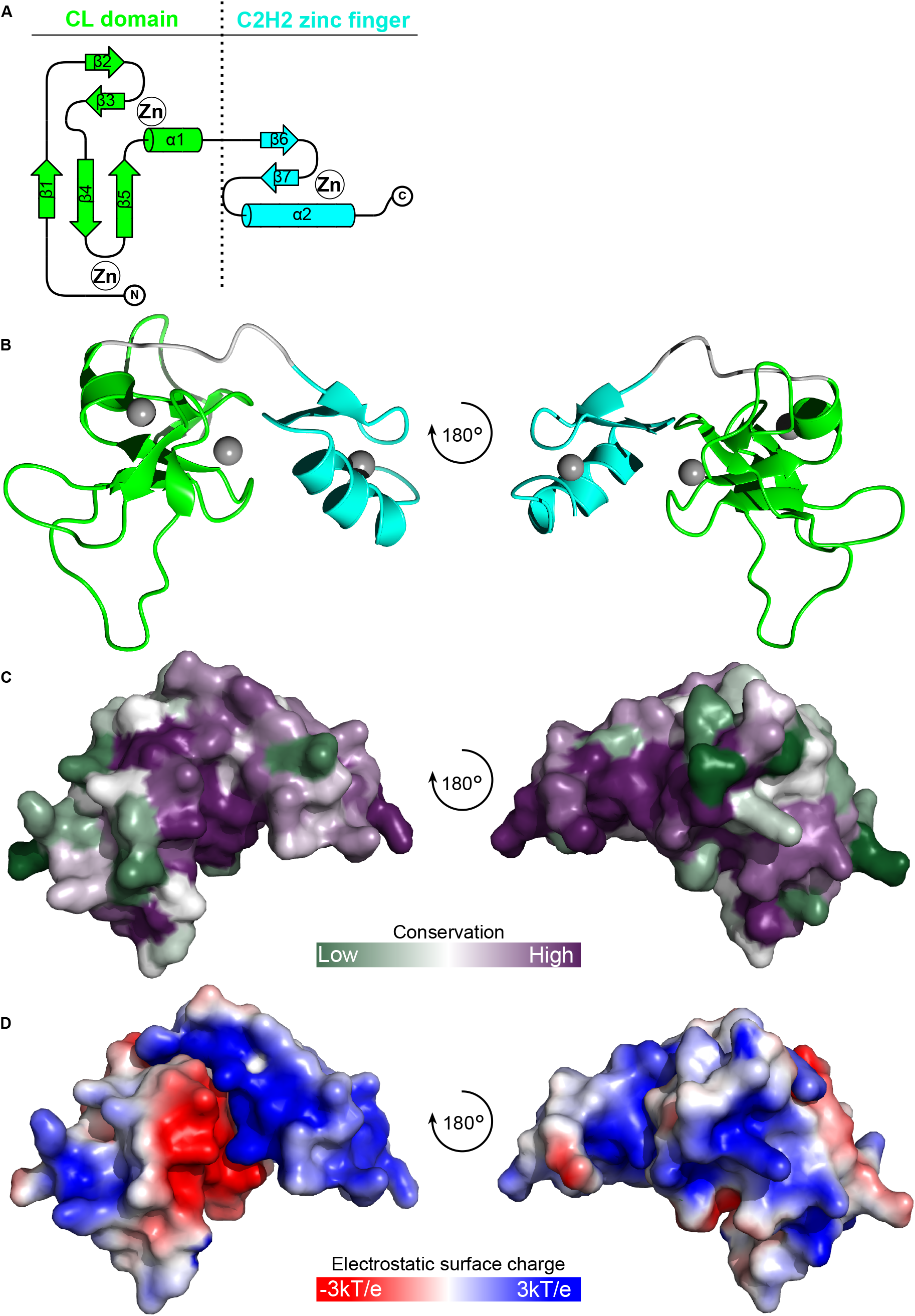
Crystal structure of *Bodo saltans* KKT2 central domain reveals the presence of two zinc-binding domains. (A) Topology diagram of the *Bs*KKT2 central domain showing the CL domain in green and C2H2-type zinc finger in cyan. (B) Cartoon representation of the *Bs*KKT2 central domain in two orientations. Zinc ions are shown in grey spheres. The structure is colored as in (A). (C) Surface representation of the *Bs*KKT2 central domain colored according to sequence conservation using the ConSurf server (Landau et al., 2005; Ashkenazy et al., 2016). Structure orientation as in (B). (D) Electrostatic surface potential of the *Bs*KKT2 central domain generated by APBS (Jurrus et al., 2018). Structure orientation as in (B).

**Table 1:**

| Data collection | *BsKKT2* central domain |
| --- | --- |
| Beamline | Diamond I04 |
| Wavelength (Å) | 1.28297 |
| Space group (Z) | I222 (8) |
| Unit cell (cell edges in Å, cell angles in degrees) | 38.96, 53.51, 83.29<br>90, 90, 90 |
| Resolution range (Å) | 45.02 – 1.8 (1.83 – 1.8) |
| Total No of reflections | 37266 (925) |
| No. of unique reflections | 8158 (334) |
| Completeness (%) | 96.96 (77.86) |
| R<sub>merge</sub> | 0.10 (0.52) |
| R<sub>pim</sub> | 0.05 (0.33) |
| CC<sub>1/2</sub> | 0.99 (0.77) |
| <I/σ(I)> | 8.65 (1.08) |
| Multiplicity | 4.57 (2.77) |
| Anomalous completeness (%) | 89.83 (37.37) |
| Anomalous multiplicity | 2.54 (1.93) |
| Overall *B* factor from Wilson plot (Å<sup>2</sup>) | 22.4 |
| Refinement |  |
| Resolution (Å) | 35.3 – 1.8 (2.0 – 1.8) |
| No. of reflections working set | 7748 (390) |
| No. of reflections test set | 403 (18) |
| Final R<sub>work</sub> (%) | 19.5 (24.3) |
| Final R<sub>free</sub> (%) | 22.5 (23.4) |
| No. of protein atoms | 770 |
| No. of Zn atoms | 3 |
| No. of water atoms | 94 |
| No. of sulfate ions | 2 |
| Average B factor (Å<sup>2</sup>) protein atoms | 30.45 |
| Average B factor (Å<sup>2</sup>) Zn atoms | 25.31 |
| Average B factor (Å<sup>2</sup>) water atoms | 44.43 |
| Rmsd bond lengths (Å) | 0.008 |
| Rmsd bond angles (°) | 0.96 |
| Ramachandran plot |  |
| Most favoured (%) | 97.89 |
| Allowed (%) | 2.11 |
| Disallowed | 0 |

In parentheses the values relative to the highest resolution shell.

A structural homology search using the DALI server (Holm and Laakso, 2016) indicated that the CL domain has weak structural similarity to proteins that have C1 domains (Table S2). C1 domains were originally discovered as lipid-binding modules in protein kinase Cs (PKCs) and are characterized by the HX_12_CX_2_CXnCX_2_CX_4_HX_2_CX_7_C motif (Colón-González and Kazanietz, 2006; Das and Rahman, 2014). C1 domains are classified into a typical C1 domain, which binds diacylglycerol or phorbol esters, and an atypical C1 domain, which is not known to bind ligands. The closest structural homolog of the *Bs*KKT2 CL domain was the atypical C1 domain of the Vav1 protein (RMSD 2.7 Å across 52 C*α*). Although the CL domain and the C1 domain share some structural similarity, their superposition revealed fundamental differences (Figure 6). Coordination of one zinc ion in the *Bs*KKT2 CL domain occurs via the N-terminal residues Cys 580 and His 584, while that in the Vav1 C1 domain occurs via the N-terminal His 516 and C-terminal Cys 564. More importantly, the CL domain does not have the HX_12_CX_2_CXnCX_2_CX_4_HX_2_CX_7_C motif that is present in all C1 domains. Therefore, the structural similarity of these two distinct domains is likely a product of convergent evolution.

**Figure 6.**
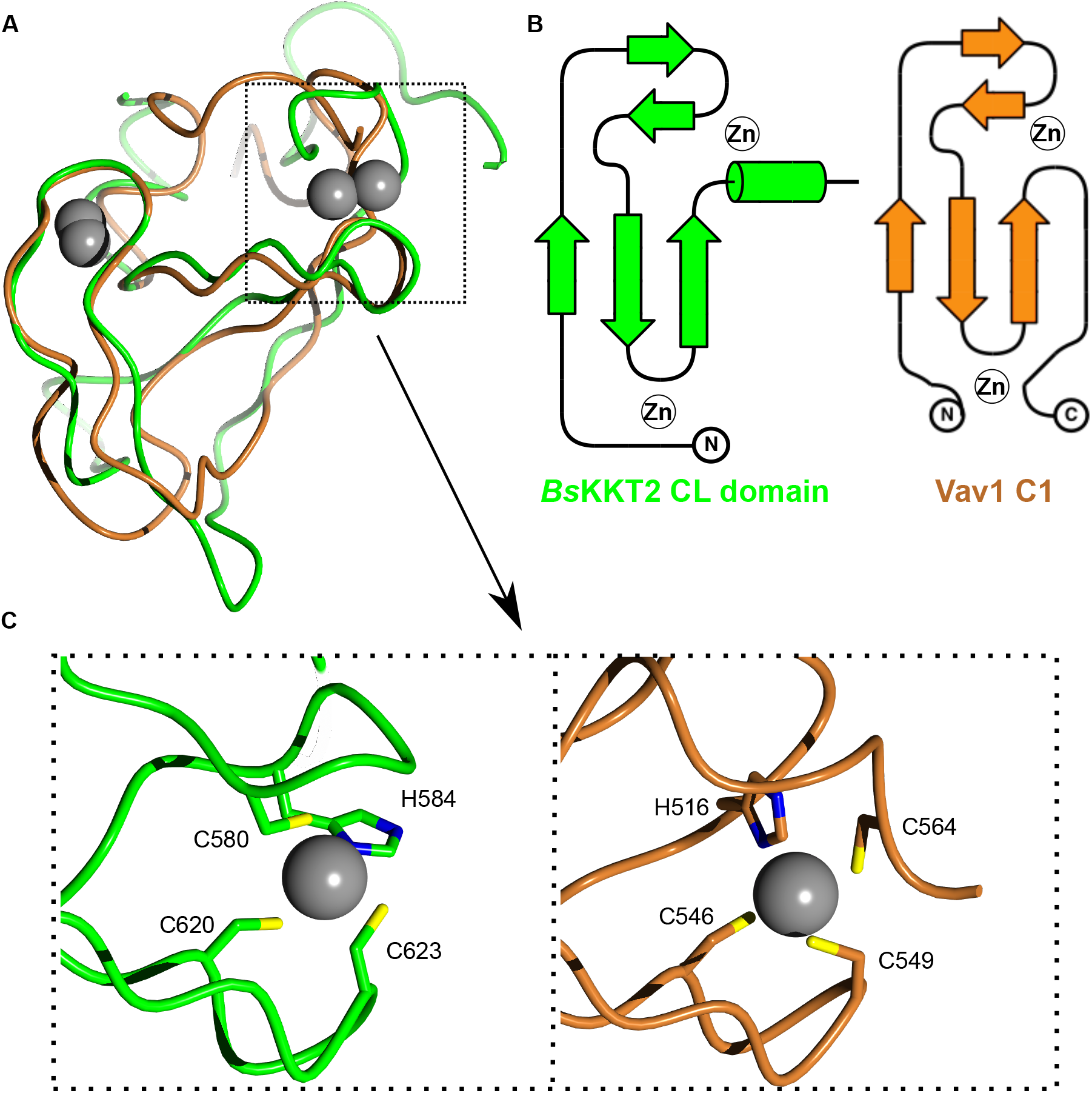
*Bodo saltans* KKT2 CL is a unique domain. (A) Structure superposition of *Bs*KKT2 CL domain in green and the Vav1 C1 domain in brown (PDB: 3KY9 (Yu et al., 2010)). Zinc ions are shown in grey spheres. (B) Topology diagram of *Bs*KKT2 CL domain and Vav1 C1 domain. (C) Close-up view showing a key difference in zinc coordination between *Bs*KKT2 CL domain and Vav1 C1 domain.

Structural analysis of the C-terminal zinc-binding domain of *Bs*KKT2 revealed a classical C2H2-type zinc finger (Table S3). C2H2 zinc fingers are known to bind DNA, RNA, or protein (Krishna et al., 2003; Brayer and Segal, 2008). In most known cases, two or more C2H2 zinc fingers are used to recognize specific DNA sequences, which is typically achieved by specific interactions between the side chain of residues in positions −1, 2, 3, and 6 in the recognition α-helix (where −1 is the residue immediately preceding the α-helix) and DNA bases (Wolfe et al., 2000). Notably, some proteins with a single zinc finger can recognize specific DNA sequences (Omichinski et al., 1997; Dathan et al., 2002). The C-terminal zinc-binding domain of *Bs*KKT2 consists of one C2H2 domain (−1: Ser 653, 2: Thr 655, 3: Lys 656, 6: Tyr 659). The sequence alignment of the *Bs*KKT2 C2H2 zinc finger shows that residues at the position −1 and 3 are highly conserved in trypanosomatids, while those at position 2 and 6 are not (Figure 8A).

### The CL domain structure is conserved in *Perkinsela* KKT2a

We next asked whether the central domain structure is conserved among kinetoplastids. *Perkinsela* is a highly divergent endosymbiotic kinetoplastid that lives inside *Paramoeba* (Tanifuji et al., 2017). Our homology search identified three proteins that have similarity to KKT2 and KKT3. Interestingly, similarities among these *Perkinsela* proteins are higher than those between them and KKT2 or KKT3 in other kinetoplastids (Figure S3). Because these *Perkinsela* proteins overall have higher sequence similarity to KKT2 than KKT3, we call them *Pk*KKT2a (XU18_4017), *Pk*KKT2b (XU18_0308), and *Pk*KKT2c (XU18_4564). *Pk*KKT2c does not have a kinase domain, like KKT20 in other kinetoplastids (Nerusheva and Akiyoshi, 2016). Our sequence alignment suggests that *Pk*KKT2a and *Pk*KKT2b have a CL-like domain but lack a C2H2 zinc finger (Figure S3).

We determined the crystal structure of *Pk*KKT2a^551–679^ at 2.9 Å resolution by Zn-SAD phasing (Figure S2 and Table 2), which confirmed the presence of a CL-like structure: 2 β-sheets (residues 551–647), followed by an extended C-terminal α-helix (residues 648–679) (Figure 7A, B). The CL-like domain of *Pk*KKT2a overlaps closely with that of *Bs*KKT2 (RMSD: 0.79 Å across 39 C*α*), with the exception of some differences being localized to the loop insertion and the absence of a C2H2 domain in *Pk*KKT2a^551–679^ (Figure 7C), consistent with our sequence analysis (Figure S3). Taken together, our structures have revealed that the CL domain is conserved in *Bs*KKT2 and *Pk*KKT2a. Given the sequence similarity of KKT2 between *Bodo saltans* and other trypanosomatids (Figure 8A), it is likely that the unique CL domain structure is conserved among kinetoplastids.

**Table 2:**

| Data collection | *PkKKT2a*<sup>551–679</sup> |  |
| --- | --- | --- |
| Beamline | Diamond I24 | Diamond I03 |
| Wavelength (Å) | 0.96861 | 1.28272 |
| Space group (Z) | P6<sub>4</sub> (6) | P6<sub>4</sub> (6) |
| Unit cell (cell edges in Å, cell angles in degrees) | 113.84, 113.84, 46.01, 90, 90, 120 | 114.33, 114.33, 46.32, 90, 90, 120 |
| Resolution range (Å) | 56.92 – 2.87 (2.92 - 2.87) | 99.01 – 3.80 (3.86 - 3.80) |
| Total No of reflections | 153076 (6844) | 68026 (3882) |
| No. of unique reflections | 7986 (357) | 3535 (189) |
| Completeness (%) | 99.73 (91.77) | 100.00 (100.00) |
| R<sub>merge</sub> | 0.10 (1.04) | 0.31 (6.13) |
| R<sub>pim</sub> | 0.024 (0.240) | 0.073 (1.38) |
| CC<sub>1/2</sub> | 0.99 (0.45) | 0.99 (0.4) |
| <I/σ(I)> | 16.45 (2.14) | 12 (1.7) |
| Multiplicity | 19.17 (19.17) | 19.2 (20.5) |
| Anomalous completeness (%) | 99.6 (91.2) | 100 (100) |
| Anomalous multiplicity | 10 (9.9) | 10.2 (10.6) |
| Overall *B* factor from Wilson plot (Å<sup>2</sup>) | 105.95 | 136.7 |
| Refinement |  |  |
| Resolution (Å) | 49 – 2.87 (3 – 2.87) |  |
| No. of reflections working set | 7408 (417) |  |
| No. of reflections test set | 377 (16) |  |
| Final R<sub>work</sub> (%) | 25.2 (36.6) |  |
| Final R<sub>free</sub> (%) | 27.5 (51.2) |  |
| No. of protein atoms | 832 |  |
| No. of Zn atoms | 2 |  |
| Average B factor (Å<sup>2</sup>) protein atoms | 113.4 |  |
| Average B factor (Å<sup>2</sup>) Zn atoms | 100.6 |  |
| Rmsd bond lengths (Å) | 0.009 |  |
| Rmsd bond angles (°) | 1.11 |  |
| Ramachandran plot |  |  |
| Most favoured (%) | 90.9 |  |
| Allowed (%) | 9.1 |  |
| Disallowed | 0 |  |

In parentheses the values relative to the highest resolution shell.

**Figure 7.**
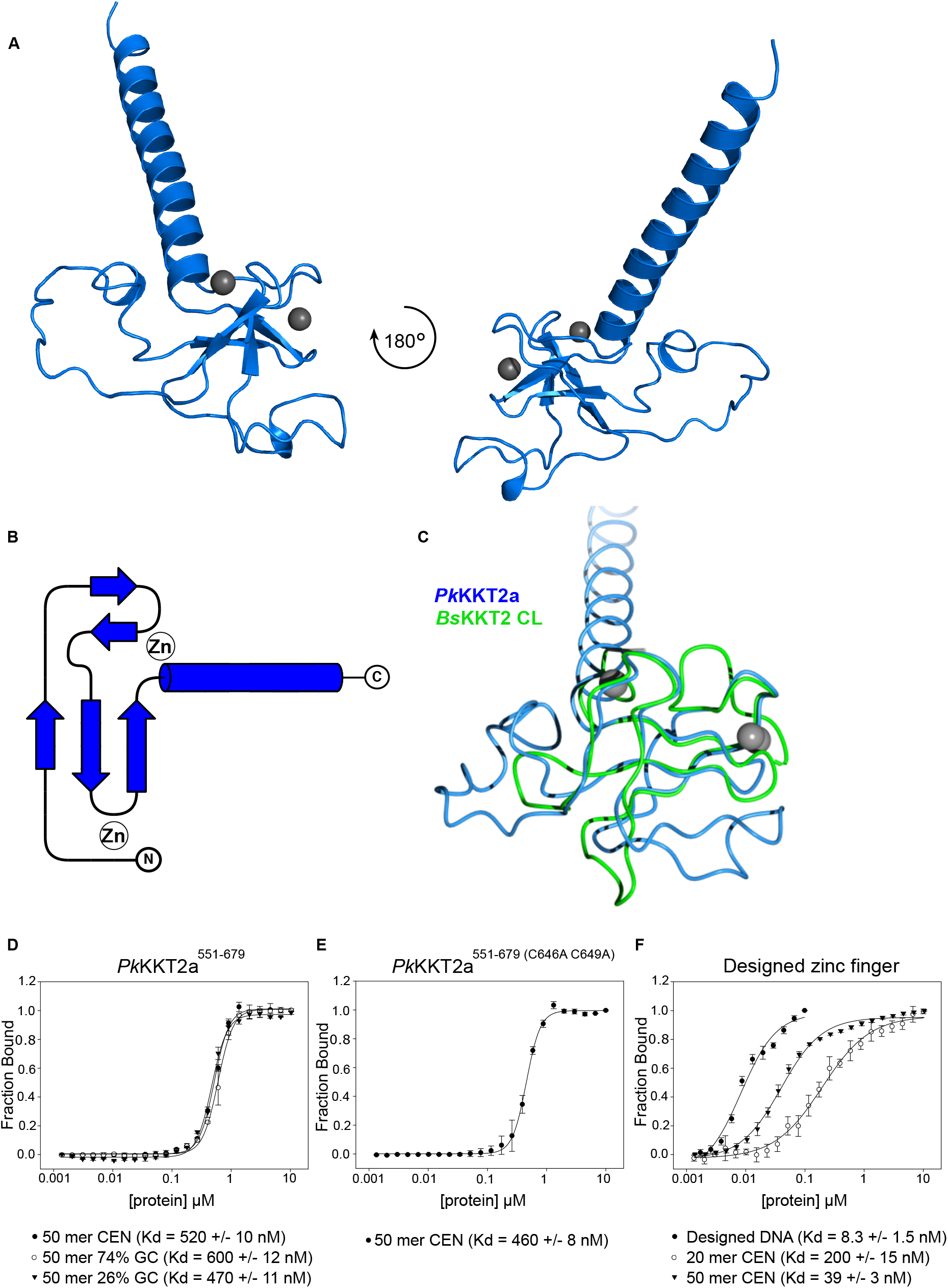
Crystal structure of *Perkinsela* KKT2a^551–679^ highlights conservation of CL domain. (A) Cartoon representation of *Pk*KKT2a^551–679^ in two orientations. Zinc ions are shown in grey spheres. (B) Topology diagram of *Pk*KKT2a^551–679^ structure. (C) Structure superposition of *Pk*KKT2a^551–679^ and *Bs*KKT2 CL domain, showing that the core of the structure is conserved. Variations between the two structures are due to sequence insertions within CL and the absence of the C2H2 zinc finger at the C-terminus in *Pk*KKT2a^551–679^. (D) Fluorescence polarization assay for *Pk*KKT2a^551–679^ on 50-bp DNA probes of different GC contents, showing that it binds DNA in a sequence-independent manner. 50 mer CEN probe is part of the centromeric sequence (CIR147) in *T. brucei* and has 36% GC content. Probes, BA1674, BA2218, BA2216. (E) Fluorescence polarization assay for *Pk*KKT2a^551–679^ ^(C646^ ^C649A)^, showing that the mutant has similar DNA-binding affinity. Probe, BA1674. (F) Fluorescence polarization assay for designed zinc finger (Jantz and Berg, 2010), showing that it has sequence-specific DNA-binding activity. 20 mer CEN DNA has 35% GC content. Probes, BA3083, BA1793, BA1674.

**Figure 8.**
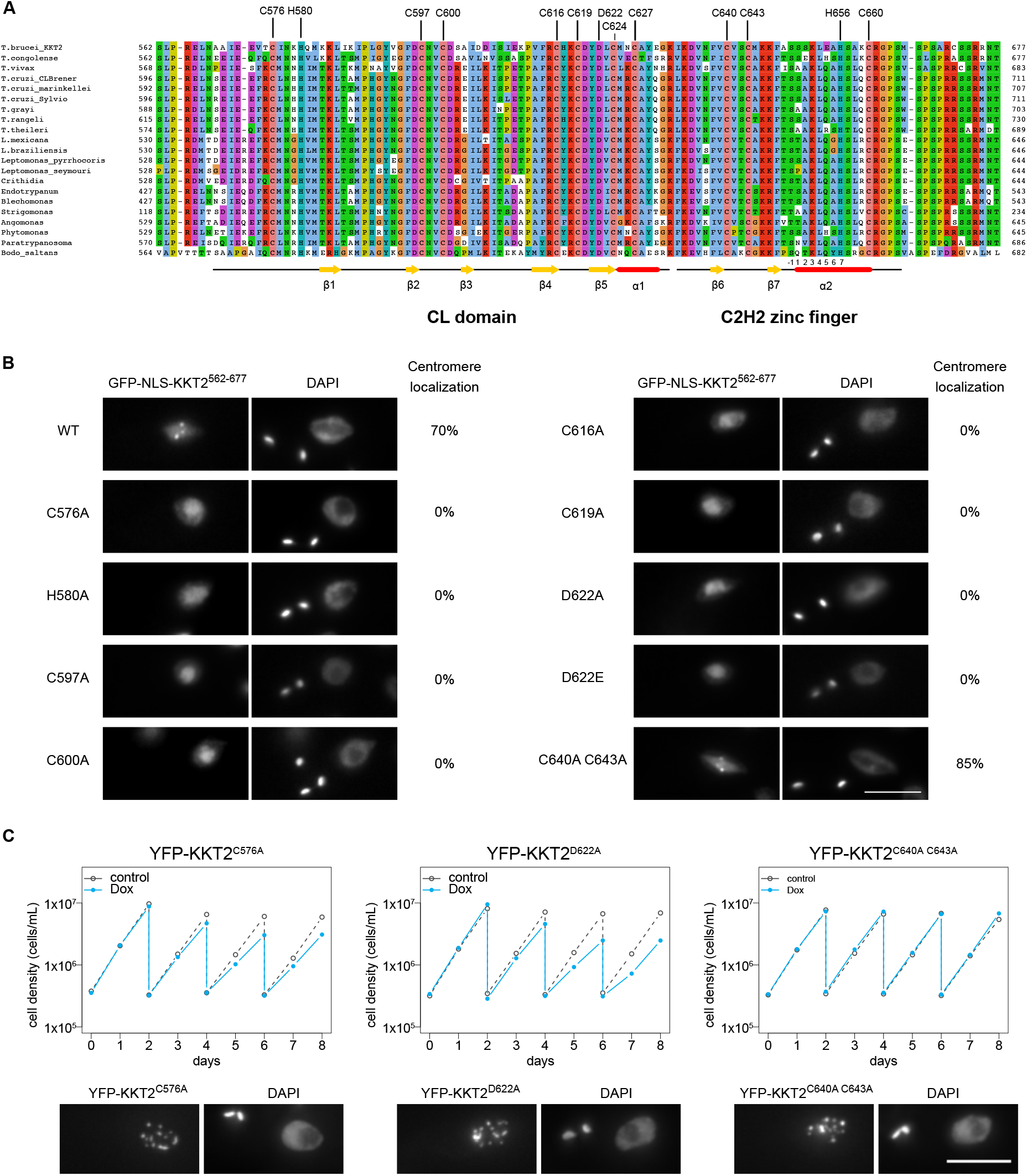
KKT2 CL domain is critical for the centromere localization in *T. brucei*. (A) Multiple sequence alignment of KKT2. Residues that coordinate zinc ions, the conserved aspartic acid residue, and secondary structures of *Bs*KKT2 are shown. Putative SPKK motifs are highlighted in boxes. Numbers below the alignment are for the C2H2 zinc finger’s α-helix, where −1 is the residue immediately preceding the helix. (B) Kinetochore localization of *Tb*KKT2^562–677^ depends on the CL domain, but not on C2H2 zinc finger. Percentages of GFP-positive 2K1N cells that have kinetochore-like dots were quantified at 1 day post-induction (n = 40, each). Inducible GFP-NLS fusion proteins were expressed with 10 ng/mL doxycycline. Cell lines, BAP457, BAP1700, BAP1702, BAP1710, BAP1712, BAP1715, BAP1717, BAP1649, BAP1719, BAP1837. (C) *Tb*KKT2 C576A and D622A mutants localize at kinetochores but fail to support normal cell growth. One allele of *Tb*KKT2 was mutated and tagged with an N-terminal YFP tag, and the other allele was depleted using RNAi-mediated knockdown by targeting the 5′UTR of the *Tb*KKT2 transcript. Top: cells were diluted every two days and cell growth was monitored for 8 days upon induction of RNAi. Controls are uninduced cell cultures. Similar results were obtained for at least three clones of *Tb*KKT2 mutants. Bottom: Example of cells expressing the *Tb*KKT2 mutants prior to RNAi induction, showing that they localize at kinetochores (n > 150, each) (also see Figure S7A). Maximum intensity projections are shown. RNAi was induced with 1 μg/mL doxycycline. Cell lines, BAP1789, BAP1779, BAP1786. Scale bars, 5 μm.

### *Perkinsela* KKT2a has DNA-binding activity

Although *Perkinsela* KKT2a lacks a C2H2 zinc finger, our sequence analysis of the KKT2 central domain revealed a putative C2H2 zinc finger not only in trypanosomatids and bodonids but also in one of Prokinetoplastina’s KKT2-like proteins, PhM_4_m.86555 (Figure S3). Furthermore, DNA-binding SPKK motifs (Suzuki, 1989) are present right after the C2H2 zinc finger in many kinetoplastids, while an AT-hook motif is present within the CL-like domain of *Perkinsela* KKT2b (Figure S3). These observations suggest that the central domain of KKT2 and KKT3 have DNA-binding activity, perhaps stabilizing their localization at centromeres. To test this hypothesis, we performed fluorescence polarization assays using fluorescently-labelled DNA probes. Unfortunately, we were unable to obtain reliable data for *T. brucei* and *B. saltans* KKT2 central domains due to fluorophore quenching. We therefore focused on *Pk*KKT2a^551–679^ that has a CL-like domain. Our fluorescence polarization assay showed that *Pk*KKT2a^551–679^ has DNA-binding activity with a *K_d_* of ~500 nM on three DNA probes that have different GC contents (50 mer CEN is 50-bp DNA sequence from the CIR147 centromere repeat in *T. brucei*) (Figure 7D). To assess the importance of the structural integrity for DNA binding, we next performed fluorescence polarization assay for *Pk*KKT2a^551–679^ that has mutations in Zn-coordinating residues (C646 and C649, which correspond to C616 and C619 in *Tb*KKT2) and found that it has a similar DNA-binding affinity compared to wild-type *Pk*KKT2a^551–679^ (Figure 7E). As a comparison, we used a well-characterized zinc finger (Designed zinc finger) that binds a specific DNA sequence (Designed DNA) (Jantz and Berg, 2010). This protein bound its optimal DNA sequence with a *K_d_* of 8 nM, while it had weaker affinity for 20-bp and 50-bp probes from the CIR147 centromere sequence (Figure 7F). These results show that *Pk*KKT2a^551–679^ has DNA-binding activity. It is possible that *Pk*KKT2a^551–679^ has higher DNA-binding activity for *Perkinsela* centromere DNA sequences (yet to be identified).

### The CL domain of KKT2 is important for long-term viability in *T. brucei*

To examine the functional relevance of the CL domain and C2H2 zinc finger, we tested their mutants in *T. brucei*. We first made various mutants in full-length *Tb*KKT2 and found that all mutants localized at kinetochores (Figure S4). Because *Tb*KKT2 has multiple domains that can independently localize at kinetochores (Figure 3), we next expressed mutants in our ectopic expression of the central domain (*Tb*KKT2^562–677^). We found that mutations in Zn-coordinating residues of the CL domain (C576A, H580A, C597A, C600A, C616A, C619A) all abolished kinetochore localization (Figure 8A, B). In contrast, similar mutations in the C2H2-type zinc finger (C640A C643A) did not affect the localization.

To gain insights into how the KKT2 CL domain may promote kinetochore localization, we analyzed conservation and electrostatic potential of the *Bs*KKT2 CL domain surface residues to identify possible patches that may be involved in this process. Our analysis revealed a highly conserved acidic patch, centered around residue *Bs*KKT2 D626 (Figure 5C, D, and Figure S5). Interestingly, this aspartic acid is strictly conserved in all KKT2 and KKT3 proteins (Figure S3). To test the importance of this residue, we mutated the corresponding residue in *T. brucei* and found that *Tb*KKT2^562–677^ with either D622A or D622E failed to localize at kinetochores (Figure 8B). Taken together, our results show that the CL domain, but not the C2H2 zinc finger, is important for the kinetochore localization of the *Tb*KKT2 central domain.

To test the importance of the *Tb*KKT2 central domain for cell viability, we next performed rescue experiments. We replaced one allele of *Tb*KKT2 with an N-terminally YFP-tagged *Tb*KKT2 construct that has either wild-type or mutant versions of the central domain, and performed RNAi against the 5′UTR of the *Tb*KKT2 transcript to knockdown the untagged allele of *Tb*KKT2 (Figure S6) (Ishii and Akiyoshi, 2020). As expected, mutants in the CL domain (C576A and D622A) and the C2H2 zinc finger (C640A C643A) both localized at kinetochores (Figure 8C and Figure S7). Upon induction of RNAi, however, the CL domain mutants failed to support normal cell growth after day 4, while the C2H2 zinc finger mutants rescued the growth defects. These data confirm the importance of the CL domain for the function of *Tb*KKT2 *in vivo*.

### Localization of KKT3 depends on the central domain in *T. brucei*

The central domain of *Tb*KKT3 can localize at kinetochores throughout the cell cycle (Figure 4). The sequence similarity of the central domain between KKT2 and KKT3 suggested that *Tb*KKT3 likely consists of two domains, which correspond to the CL domain and the C2H2 zinc finger present in KKT2 (Figure S3). Consistent with this prediction, mutating *Tb*KKT3 residues that align with Zn-coordinating histidine or cysteine residues in the CL domain of KKT2 abolished the kinetochore localization of the ectopically-expressed full length *Tb*KKT3 protein (Figure 9A, B). We also found that the conserved aspartic acid D692 was essential for kinetochore localization. In contrast, mutations in the *Tb*KKT3 C2H2 zinc finger (C707A C710A) did not affect the kinetochore localization.

**Figure 9.**
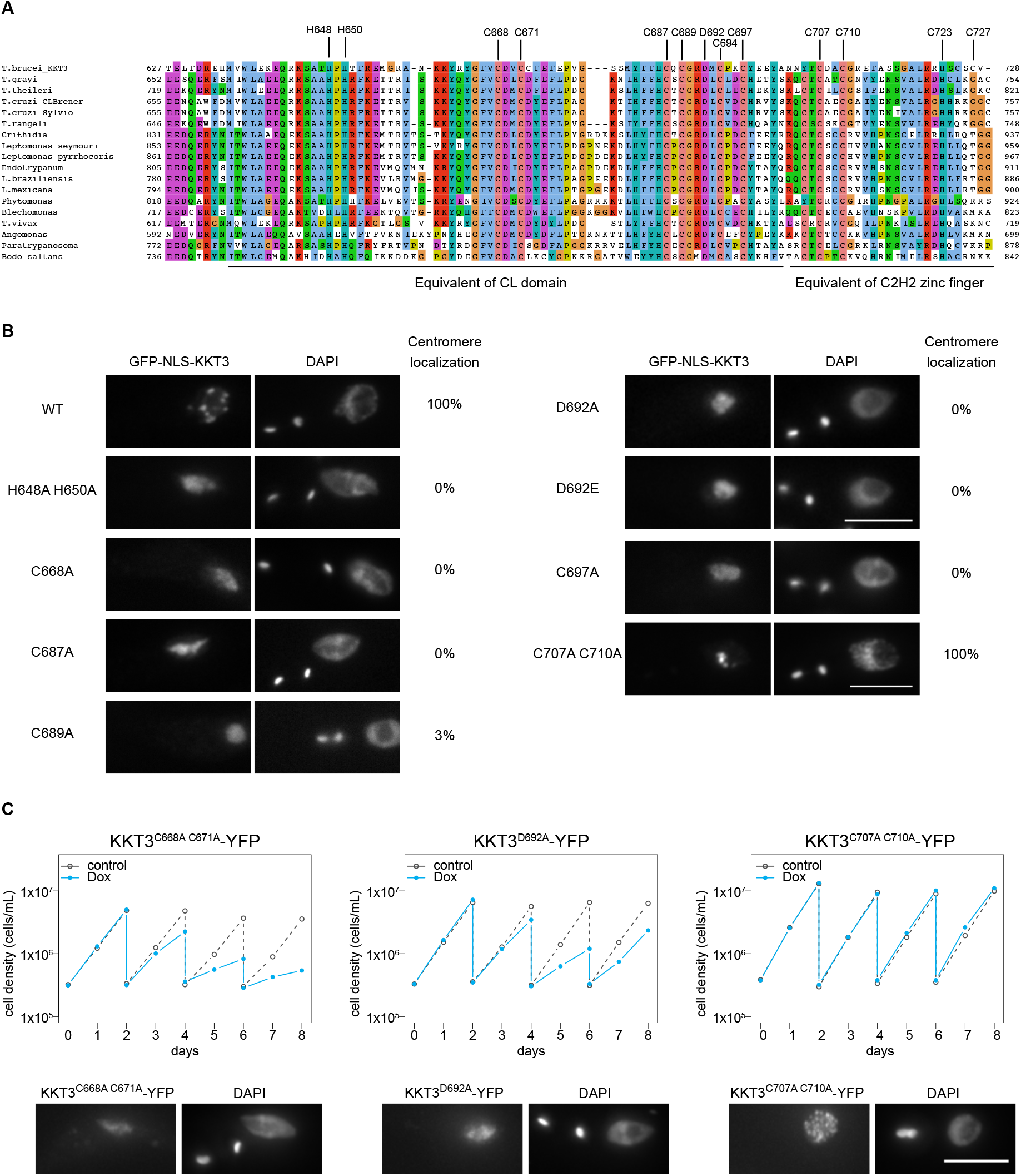
Kinetochore localization of KKT3 depends on the central domain in *T. brucei*. (A) Multiple sequence alignment of KKT3. Residues that are expected to coordinate zinc ions as well as the conserved aspartic acid residue are shown. (B) Percentage of GFP-positive cells that have kinetochore-like dots were quantified at 1 day post-induction (n > 22, each). Inducible GFP-NLS fusion proteins were expressed with 10 ng/mL doxycycline. Cell lines, BAP291, BAP359, BAP360, BAP446, BAP447, BAP1721, BAP1722, BAP362, BAP341. (C) *Tb*KKT3 C668A/C671A and D692A mutants do not localize at kinetochores and fail to support normal cell growth, while *Tb*KKT3 C707A/C710A mutant is functional. One allele of *Tb*KKT3 was mutated and tagged with a C-terminal YFP tag, and the other allele was depleted using RNAi-mediated knockdown by targeting the 3′UTR of the *Tb*KKT3 transcript. Top: cells were diluted every two days and cell growth was monitored for 8 days upon induction of RNAi. Similar results were obtained for at least three clones of *Tb*KKT3 mutants. Controls are uninduced cell cultures. Bottom: Example of cells expressing the *Tb*KKT3 mutants prior to RNAi induction, showing that *Tb*KKT3^C668A^ ^C671A^ and *Tb*KKT3^D692A^ do not localize at kinetochores while *Tb*KKT3^C707A^ ^C710A^ localizes normally (n > 90, each) (also see Figure S7B). Maximum intensity projections are shown. RNAi was induced with 1 μg/mL doxycycline. Cell lines, BAP1791, BAP1793, BAP1783. Scale bars, 5 μm.

We next performed rescue experiments by replacing one allele of *Tb*KKT3 with a C-terminally YFP-tagged construct that has either wild-type or mutant versions of the central domain, and performed RNAi against the 3′UTR of *Tb*KKT3 to knockdown the untagged allele of *Tb*KKT3 (Figure S6). We first confirmed that *Tb*KKT3 CL domain mutants (C668A C671A and D692A) were unable to localize at kinetochores, while the *Tb*KKT3 C2H2 zinc finger mutant (C707A C710A) localized normally (Figure 9C and Figure S7). Upon induction of RNAi, CL mutants failed to rescue the growth defect, showing that kinetochore localization is essential for the *Tb*KKT3 function (Figure 9C). In contrast, the *Tb*KKT3 C2H2 zinc finger mutant supported normal cell growth. These data show that the *Tb*KKT3 CL domain is essential for the localization and function of *Tb*KKT3.

## Discussion

A major open question concerning the biology of kinetoplastids is how these organisms assemble kinetochores specifically at centromeres using a unique set of kinetochore proteins. Studies in other eukaryotes have shown that constitutively localized kinetochore proteins (such as CENP-A and CENP-C) play crucial roles in kinetochore specification and assembly (French and Straight, 2017; Hamilton and Davis, 2020; Kixmoeller et al., 2020). Among the six proteins that constitutively localize at kinetochores in *T. brucei* (KKT2, KKT3, KKT4, KKT20, KKT22, and KKT23), we previously showed that KKT4 is important for recruiting KKT20 but not many other proteins including KKT2 and KKT3 (Llauró et al., 2018). In this study, we show that KKT2 and KKT3 are important for recruiting multiple kinetochore proteins (including KKT4, KKT22, and KKT23), while their localization is independent from various kinetochore proteins. Together with the fact that KKT2 and KKT3 have DNA-binding motifs, these results support a hypothesis that KKT2 and KKT3 locate at the base of kinetoplastid kinetochores and play crucial roles in recruiting other kinetochore proteins. It will be important to identify which proteins directly interact with KKT2/3 to understand the mechanism of kinetochore assembly.

KKT2/3 have sequence similarity with polo-like kinases (Nerusheva and Akiyoshi, 2016). In addition to an N-terminal protein kinase domain and a C-terminal divergent polo boxes, they have a central domain that is highly conserved among kinetoplastids. By ectopically expressing fragments of KKT2 and KKT3 in *T. brucei*, we established that their central domains can localize specifically at centromeres. The crystal structure of the *Bodo saltans* KKT2 central domain revealed a unique structure, which consists of two distinct zinc-binding domains, the CL domain and C2H2-type zinc finger. It is likely that the central domain of *T. brucei* KKT2 has a similar structure based on high sequence similarity between *Bs*KKT2 and *Tb*KKT2 proteins. Importantly, mutational analyses of *Tb*KKT2 revealed that the CL domain is important for the localization of the central domain, while the C2H2 zinc finger is not. Furthermore, although full-length *Tb*KKT2 CL mutants localized at kinetochores (likely due to interactions with other kinetochore proteins via other domains of *Tb*KKT2), they were not fully functional. Taken together, these data have established that the CL domain is essential for the function of *Tb*KKT2, which is consistent with the presence of CL, but not C2H2 zinc finger, in *Perkinsela* KKT2a. It remains unclear whether the structure of the central domain is conserved between KKT2 and KKT3. Nonetheless, our functional studies showed that the equivalent domain of CL in *Tb*KKT3 was also essential for the kinetochore localization and function, while the equivalent domain of C2H2 zinc finger was not, showing that the functional importance of CL is conserved in KKT3. It will be important to obtain KKT3 central domain structures to reveal structural similarity or difference between KKT2 and KKT3. It is noteworthy that all identified KKT2/3 homologs in deep-branching Prokinetoplastina have higher similarity to KKT2 than KKT3. We speculate that ancestral kinetoplastids had only KKT2-like proteins that carried out all necessary functions and that KKT3 in trypanosomatids and bodonids represents a product of gene duplication that became specialized in certain functions (such as more efficient centromere localization by its central domain compared to KKT2).

It remains unclear how kinetoplastids specify kinetochore positions. The fact that KKT2/3 central domains manage to localize at centromeres suggests that they are able to recognize something special at centromeres. What might be a unique feature at centromeres in kinetoplastids that lack CENP-A? Histone variants are one possibility. *T. brucei* has four histone variants, H2AZ, H2BV, H3V, and H4V. However, none of them is specifically enriched at centromeres (Lowell and Cross, 2004; Lowell et al., 2005; Siegel et al., 2009), and histone chaperones did not co-purify with any kinetochore protein (Akiyoshi and Gull, 2014). Alternatively, there might exist certain post-translational modifications on histones or DNA specifically at centromeres (e.g. phosphorylation, methylation, acetylation, ubiquitination, or sumoylation). The KKT2 CL domain has a highly conserved acidic patch, which might act as a “reader” for such modifications. Although there is no known histone or DNA modification that occurs specifically at centromeres, KKT2/3 have a protein kinase domain and KKT23 has a Gcn5-related N-acetyltransferase (GNAT) domain (Nerusheva et al., 2019). It will be important to examine whether these enzymatic domains are important for proper recruitment of KKT2/3 central domains. Another unique feature at centromeres is the presence of kinetochore proteins, which could potentially recruit newly synthesized kinetochore components by direct protein-protein interactions. Finally, it is important to note that it remains unclear whether kinetoplastid kinetochores build upon nucleosomes. It is formally possible that the KKT2/3 central domains directly bind DNA and form a unique environment at centromeres. Understanding how the KKT2/3 central domains localize specifically at centromeres will be key to elucidating the mechanism of how kinetoplastids specify kinetochore positions in the absence of CENP-A.

## Materials and Methods

### Tryp cells and plasmids, microscopy, immunoprecipitation, and mass spectrometry

All trypanosome cell lines, plasmids, primers, and synthetic DNA used in this study are listed in Table S4. All trypanosome cell lines used in this study were derived from *T. brucei* SmOxP927 procyclic form cells (TREU 927/4 expressing T7 RNA polymerase and the tetracycline repressor to allow inducible expression) (Poon et al., 2012). Cells were grown at 28°C in SDM-79 medium supplemented with 10% (v/v) heat-inactivated fetal calf serum (Brun and Schönenberger, 1979). Endogenous YFP tagging was performed using the pEnT5-Y vector (Kelly et al., 2007). Endogenous tdTomato tagging was performed using pBA148 (Akiyoshi and Gull, 2014) and its derivatives. Inducible expression of GFP-NLS fusion proteins was performed using pBA310 (Nerusheva and Akiyoshi, 2016).

To make pBA1711 (KKT3 3′UTR hairpin RNAi construct targeting the KKT3 transcript from stop codon to +370 bp), BAG95 synthetic DNA fragment was digested with HindIII/BamHI and subcloned into the HindIII/BamHI sites of pBA310. pBA2052 (KKT2/3 double hairpin RNAi construct targeting KKT2 5′UTR from −342 bp to start codon as well as KKT3 3′UTR from stop codon to +370 bp) was made with BAG129 as above. Similarly, pBA861 (KKT1 hairpin RNAi targeting 1351–1776 bp) was made with BAG24, pBA864 (KKT6 hairpin RNAi targeting 12–499 bp) with BAG27, pBA869 (KKT14 hairpin RNAi targeting 859–1274 bp) with BAG32, pBA1316 (KKT8 3′UTR hairpin RNAi targeting from +31 to +446 bp) with BAG79, pBA1845 (KKT22 hairpin RNAi targeting 466 bp to stop codon) with BAG105, pBA1997 (KKT24 hairpin RNAi targeting 1001–1400 bp) with BAG115, and pBA2021 (KKT23 hairpin RNAi targeting 635–1047 bp) with BAG127. To make pBA1091 (KKIP1 RNAi), 297–856 bp of KKIP1 coding sequence was amplified with primers BA1541/BA1543 and cloned into p2T7-177 using BamHI/HindIII sites (Wickstead et al., 2002). To make pBA1807 (C-terminal YFP-tagging of KKT3), 4–3174 bp of KKT3 coding sequence and 250 bp of 3′UTR were amplified with primers BA2351/BA2352 and BA2353/BA2354, digested with HindIII/NotI and NotI/SpeI, respectively, and were cloned into the pEnT5-Y using HindIII/SpeI sites. Details of other plasmids are described in Table S4. Site-directed mutagenesis was performed using primers and template plasmids listed in Table S4. All constructs were sequence verified.

Plasmids linearized by NotI were transfected into trypanosomes by electroporation into an endogenous locus (pEnT5-Y derivatives and pBA148/pBA192/pBA892 derivatives) or 177 bp repeats on minichromosomes (pBA310 derivatives and p2T7-177 derivatives). Concentrations of drugs used were as follows: 5 μg/ml for phleomycin, 25 μg/ml for hygromycin, 10μg/ml for blasticidin, 30 μg/ml for G418, and 1 μg/ml for puromycin. To obtain endogenously-tagged clonal strains, transfected cells were selected by the addition of appropriate drugs and cloned by dispensing dilutions into 96-well plates. Clones that express mutant versions of KKT2 or KKT3 from the endogenous locus were screened by Sanger sequencing of genomic DNA. Expression of GFP-NLS fusion proteins (pBA310 derivatives) was induced by the addition of doxycycline (10 ng/ml). RNAi was induced by the addition of doxycycline (1 μg/ml).

Fluorescence microscopy, immunoprecipitation of GFP-fusion proteins, and mass spectrometry were performed essentially as described previously (Nerusheva and Akiyoshi, 2016; Ishii and Akiyoshi, 2020). Proteins identified with at least two peptides were considered as significant and shown in Table S1.

### Multiple sequence alignment

Protein sequences and accession numbers for KKT2 and KKT3 homologs were retrieved from TriTryp database (Aslett et al., 2010), Wellcome Sanger Institute (https://www.sanger.ac.uk/), UniProt (UniProt Consortium, 2019), or a published study (Butenko et al., 2020). Searches for KKT2/3 homologs in Prokinetoplastina and Bodonida were done using hmmsearch on its predicted proteome using manually prepared KKT2/3 hmm profiles (HMMER version 3.0 (Eddy, 1998)). Multiple sequence alignment was performed with MAFFT (L-INS-i method, version 7) (Katoh et al., 2019) and visualized with the Clustalx coloring scheme in Jalview (version 2.10) (Waterhouse et al., 2009).

### Protein expression and purification

Multiple sequence alignment together with secondary structure predictions of the KKT2 central domain were used to design constructs in *Bodo saltans* and *Perkinsela*. To make pBA1660 (*Bs*KKT2^572–668^ with an N-terminal TEV-cleavable hexahistidine (His6) tag), the central domain of *Bodo saltans* KKT2 (accession number BSAL_50690) was amplified from BAG50 (a synthetic DNA that encodes *Bodo saltans* KKT2, codon optimized for expression in Sf9 insect cells (Table S4)) with primers BA2117/BA2118 and cloned into the RSFDuet-1 vector (Novagen) using BamHI/EcoRI sites with an NEBuilder HiFi DNA Assembly Cloning Kit (NEB) according to the manufacturer’s instructions. To make pBA1139 (His6-*Pk*KKT2^551–679^), the central domain of *Perkinsela* CCAP 1560/4 KKT2a (accession number XU18_4017) was amplified from BAG48 (a synthetic DNA that encodes *Perkinsela* KKT2a, codon optimized for expression in Sf9 insect cells (Table S4)) with primers BA1569/BA1570 and cloned into RSFDuet-1 using BamHI/EcoRI sites with an In-Fusion HD Cloning Plus kit (Takara Bio). To make pBA2276 (His6-Designed zinc finger), Designed zinc finger domain was amplified from BAG136 (a synthetic DNA codon optimized for expression in *E. coli*) with primers BA3077/BA3078 and cloned into RSFDuet-1 using BamHI/EcoRI sites with an NEBuilder HiFi DNA Assembly Cloning Kit. Recombinant proteins were expressed in BL21(DE3) *E. coli* cells at 20°C using auto induction media (Formedium) (Studier, 2005).

Briefly, 500 mL of cells were grown at 37°C in 2.5 L flasks at 300 rpm until OD600 of 0.2–0.3 and then cooled down to 20°C overnight (2 L for *Bs*KKT2, 6 L for *Pk*KKT2a, and 2L for Designed zinc finger). Cells were harvested by centrifugation and resuspended in 50 ml per litre of culture of lysis buffer (25 mM Hepes pH 7.5, 150 mM NaCl, 1 mM TCEP, 10 mM imidazole, and 1.2 mM PMSF). Proteins were extracted by mechanical cell disruption using a French press (1 passage at 20,000 PSI) and the resulting lysate was centrifuged at 48,384 g for 30 min at 4°C. Clarified lysate was incubated with 5 mL TALON beads (Takara Bio), washed with 150 mL lysis buffer and eluted in 22 mL of elution buffer (25 mM Hepes pH 7.5, 150 mM NaCl, 1 mM TCEP, and 250 mM imidazole) in a gravity column, followed by TEV treatment for the removal of the His6 tag. Salt concentration of the sample was subsequently reduced to 50 mM NaCl using buffer A (50 mM Hepes pH 7.5, and 1 mM TCEP) and the sample was loaded onto a 5 mL HiTrap Heparin HP affinity column (GE healthcare) pre-equilibrated with 5% buffer B (50 mM Hepes pH 7.5, 1 M NaCl, and 1 mM TCEP) on an ÄKTA pure 25 system. Protein was eluted by using a gradient from 0.05 to 1 M NaCl, and protein-containing fractions were combined, concentrated with an Amicon stirred cell using an ultrafiltration disc with 10 kDa cut-off (Merck), and then loaded onto a HiPrep Superdex 75 16/60 size exclusion chromatography column (GE healthcare) pre-equilibrated with 25 mM Hepes pH 7.5, 150 mM NaCl, and 1 mM TCEP. Fractions containing the protein of interest were pooled together, concentrated with an Amicon stirred cell using an ultrafiltration disc with 10 kDa cut-off, and stored at −80°C. Designed zinc finger was buffered exchanged into 50 mM Tris pH 7.5, 1 mM ZnCl_2_, 50 mM NaCl and 1 mM TCEP prior storage. Protein concentration was measured by Bradford assay.

### Crystallization

Both *Bs*KKT2^572–668^ and *Pk*KKT2a^551–679^ crystals were optimized at 4°C in sitting drop vapour diffusion experiments in 48-well plates, using drops of overall volume 400 nL, mixing protein and mother liquor in a 3:1 protein:mother liquor ratio. *Bs*KKT2 central domain crystals grew from the protein at 26 mg/mL and mother liquor 40% PEG 400, 0.2 mM (NH4)2SO4, and 100 mM Tris-HCl pH 8. The 40% PEG400 in the mother liquor served as the cryoprotectant when flash-cooling the crystals by plunging into liquid nitrogen. *Pk*KKT2a^551–679^ crystals grew from the protein at 13 mg/mL and mother liquor 19% MPD, 50 mM Hepes pH 7.5, and 10 mM MgCl_2_. The crystals were briefly transferred into a cryoprotecting solution of 30% MPD, 50 mM Hepes pH 7.5, 10 mM MgCl_2_ prior to flash-cooling.

### Data collection and structure determination

X-ray diffraction data from a *Bs*KKT2 central domain crystal were collected at the I04 beamline at the Diamond Light Source (Harwell, UK) at the Zinc K-edge wavelength (λ=1.28297 Å). A set of 1441 images were processed in space group I222 using the Xia2 pipeline (Winter, 2010), with DIALS for indexing and integration (Winter et al., 2018) and AIMLESS for scaling (Evans and Murshudov, 2013) to 1.8 Å resolution. Initial 3 Zn atoms were localized by interpreting the anomalous difference Patterson, single anomalous dispersion (SAD) phases were estimated using Crank2 (Skubák and Pannu, 2013), and an initial model was built with BUCCANEER (Cowtan, 2006). The structure was completed by several cycles of alternating model building in Coot (Emsley et al., 2010) and refinement in autoBUSTER (Blanc et al., 2004; Bricogne et al., 2017).

*Pk*KKT2a^551–679^ X-ray diffraction data were collected at the I03 beamline at Diamond Light Source (Harwell, UK) also at the zinc K-edge (λ=1.28272 Å) and processed using the autoPROC pipeline (Vonrhein et al., 2011) using XDS (Kabsch, 2010) for indexing/integration and AIMLESS (Evans and Murshudov, 2013) for scaling, to a resolution of 3.8 Å. Two initial Zn positions were determined by interpreting the anomalous difference Patterson, and single anomalous dispersion (SAD) phases were estimated using Crank2 (Skubák and Pannu, 2013) and SHARP (Vonrhein et al., 2007) in space group P64. An initial model was manually built in Coot and refined once with RosettaMR (Terwilliger et al., 2012). The structure was completed by several cycles of alternating model building in Coot (Emsley et al., 2010) and refinement in autoBUSTER (Blanc et al., 2004; Bricogne et al., 2017).

A higher resolution dataset was collected from a *Pk*KKT2a^551–679^ crystal at the I24 beamline at Diamond Light Source (Harwell, UK), at a wavelength of λ=0.9686 Å. Data were processed using Xia2 pipeline (Winter, 2010), DIALS (Winter et al., 2018), and AIMLESS (Evans and Murshudov, 2013) in space group P64 to a resolution of 2.9 Å. The model obtained from the 3.8 Å dataset was used for further model building and refinement with autoBuster (Bricogne et al., 2017) and Coot (Emsley et al., 2010).

All the images were made with Pymol (Schrödinger LLC, Portland, OR) and CCP4mg (McNicholas et al., 2011). Topology diagrams were generated using TopDraw (Bond, 2003). Protein coordinates have been deposited in the RCSB Protein Data Bank (http://www.rcsb.org/) with accession codes 6TLY (*Bodo saltans* KKT2) and 6TLX (*Perkinsela* KKT2a).

### Fluorescent anisotropy DNA-binding assay

All experiments were performed in binding buffer (25 mM Hepes pH 7.5, 50 mM NaCl and 1 mM TCEP) using 1 nM FAM-labelled dsDNA sequences purchased from IDT (Table S4). Prior to the assay, proteins were buffer exchanged into binding buffer using a Zeba spin desalting column (Thermo Fisher), serially diluted at 2:3 v/v ratio and then incubated with DNA for 20 min at room temperature. Fluorescence anisotropy was measured at 25°C using a PHERAstar FS next-generation microplate reader (BMG-Labtech). Each data point is an average of 3 independent experiments. Data were fitted with SigmaPlot using a standard four-parameter logistic equation to calculate *K_d_*.

## Acknowledgments

We thank Pietro Roversi and Matt Higgins for advice, crystallography facility manager Edward Lowe, Micron Advanced Bioimaging Unit, and Advanced Proteomics Facility. We also thank Pietro Roversi, Matt Higgins, Danny Huang, and Patryk Ludzia for comments on the manuscript. Midori Ishii was supported by a long-term fellowship from the TOYOBO Biotechnology Foundation. Bungo Akiyoshi was supported by a Wellcome Trust Senior Research Fellowship (grant no. 210622/Z/18/Z) and the European Molecular Biology Organization Young Investigator Program. The authors declare that no competing interests exist.

## Author contributions

Gabriele Marcianò purified recombinant proteins, solved crystal structures, and performed DNA-binding assays. Midori Ishii performed RNAi and rescue experiments. Olga Nerusheva expressed KKT2 and KKT3 truncations and mutants in trypanosomes and performed immunoprecipitation and mass spectrometry. Bungo Akiyoshi performed RNAi experiments and expressed KKT2 and KKT3 truncations and mutants in trypanosomes. Gabriele Marcianò, Midori Ishii, and Bungo Akiyoshi wrote the manuscript.

## Data availability

All data generated during this study are included in the manuscript and supplementary data. Protein coordinates have been deposited in the RCSB Protein Data Bank (http://www.rcsb.org/x) with accession codes 6TLY (*Bodo saltans* KKT2) and 6TLX (*Perkinsela* KKT2a).

## Supplemental Tables (Excel files)

**Table S1: List of proteins identified in the immunoprecipitates of YFP-tagged KKT2^558–679^, KKT2^672–1030^, KKT2^1024–1260^, KKT2^1024–1260 (W1048A)^, KKT3^594–811^, and KKT3^846–1058^ by mass spectrometry**

**Table S2: List of DALI search hits for the***Bs***KKT2 CL domain structure**

**Table S3: List of DALI search hits for the***Bs***KKT2 C2H2 zinc finger structure**

**Table S4: List of trypanosome cell lines, plasmids, primers, synthetic DNA, and DNA probes used in this study**

## Supplementary Figure legends

**Figure S1.**
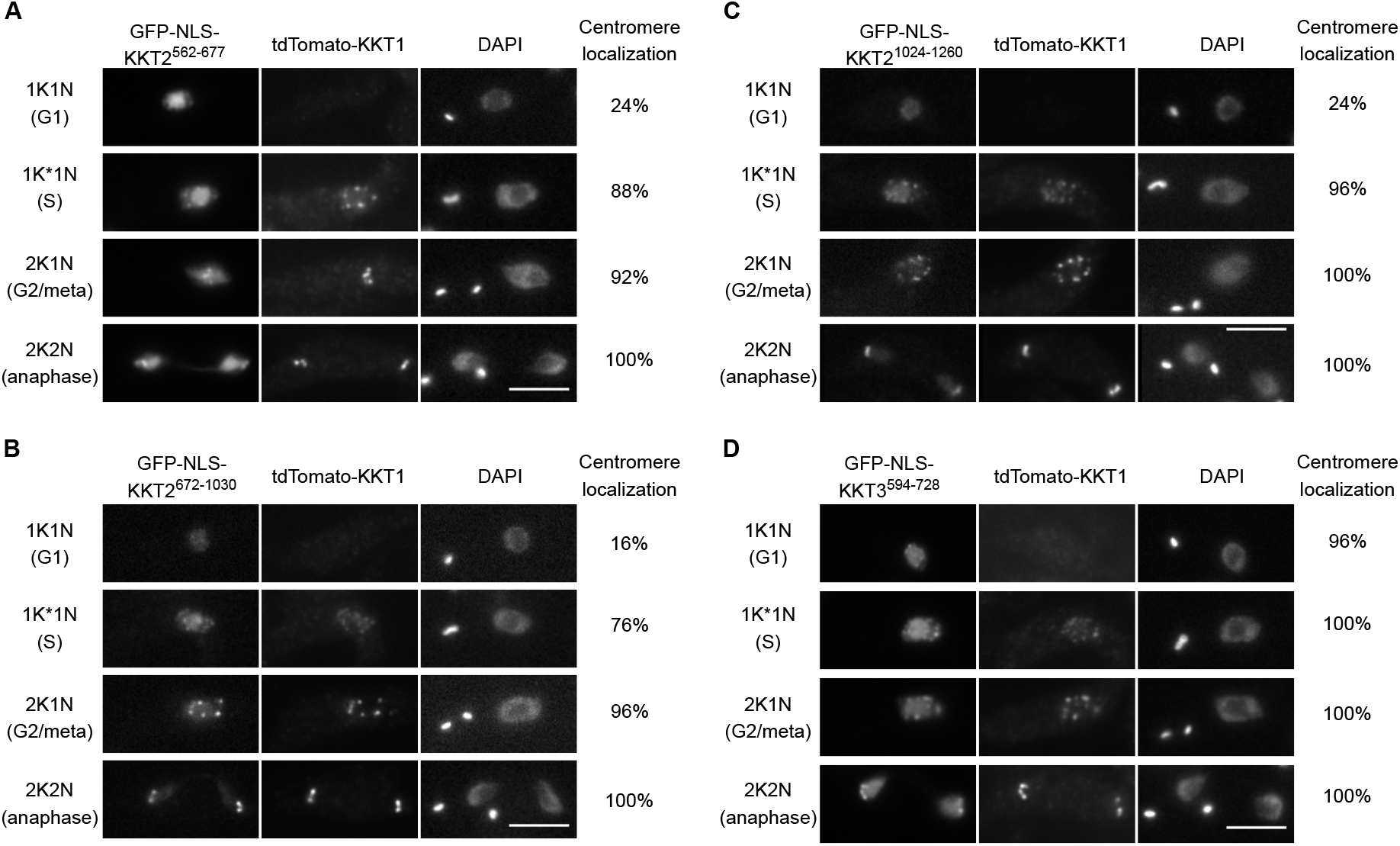
KKT2 fragments localize at kinetochores from S phase to anaphase, while KKT3 central domain localizes constitutively. (A–C) Ectopically expressed *Tb*KKT2 central domain (562–677), *Tb*KKT2^672–1030^, and *Tb*KKT2 DPB (1024–1260) localize at kinetochores from S phase until anaphase. (D) Ectopically expressed *Tb*KKT3 central domain (594–728) forms kinetochore-like dots throughout the cell cycle. Inducible GFP-NLS fusion proteins were expressed with 10 ng/mL doxycycline. tdTomato-KKT1 was used as a kinetochore marker. Cell lines, BAP1998, BAP2000, BAP1999, BAP1997. Scale bars, 5 μm.

**Figure S2.**
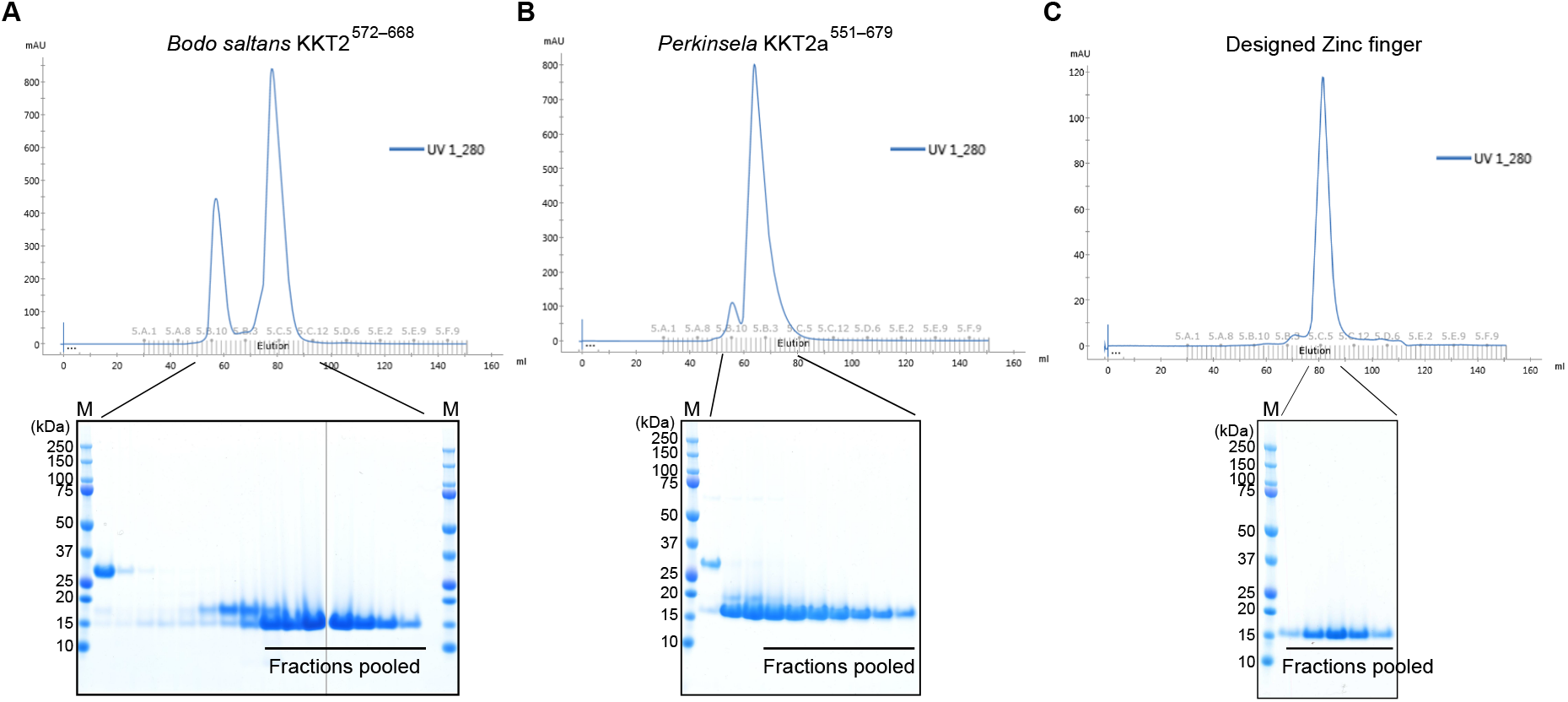
Purification of *Bodo saltans* KKT2, *Perkinsela* KKT2a, and designed zinc finger proteins. Size exclusion chromatography of (A) *Bodo saltans* KKT2 central domain, (B) *Pk*KKT2a^551–679^, and (C) designed zinc finger with respective SDS-PAGE gels showing pooled fractions. HiPrep Superdex 75 16/60 column was used. Note that two separate gels are shown for (A). M stands for protein marker.

**Figure S3.**
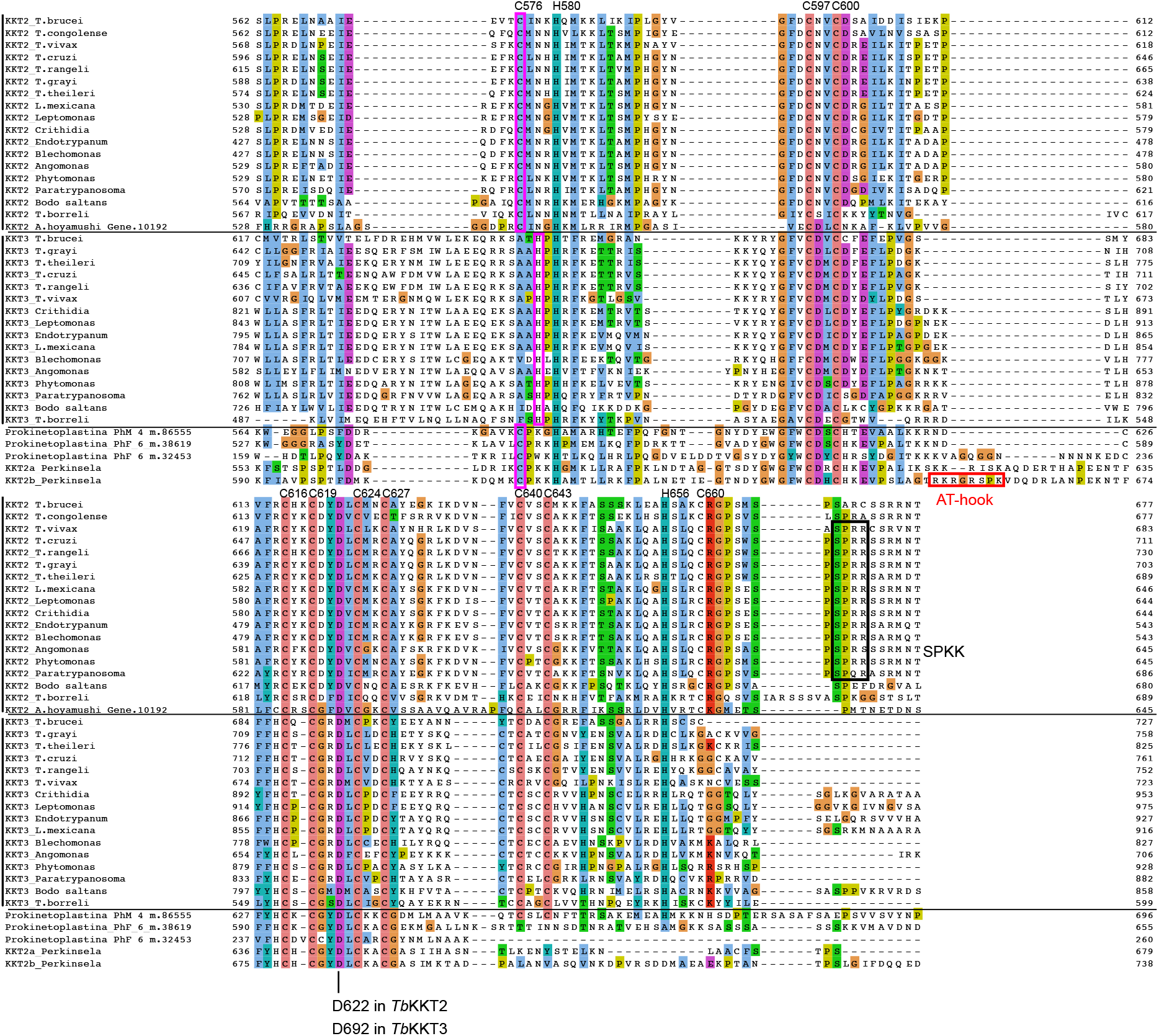
Multiple sequence alignment of KKT2 and KKT3 homologs in kinetoplastids. Residue numbers in *Tb*KKT2 for those cysteines and histidines that coordinate zinc ions in the *Bs*KKT2 structure are listed at the top of the alignment to highlight the conservation of these residues among kinetoplastids. Note that cysteine is used in KKT2 and KKT2-like proteins (C576 in *Tb*KKT2), while histidine is present in KKT3 in slightly different position (highlighted in pink box). Position of the strictly conserved aspartic acid residue is also shown. SPKK motifs (black box) and a putative AT-hook motif in *Perkinsela* KKT2b (red box) are also highlighted.

**Figure S4.**
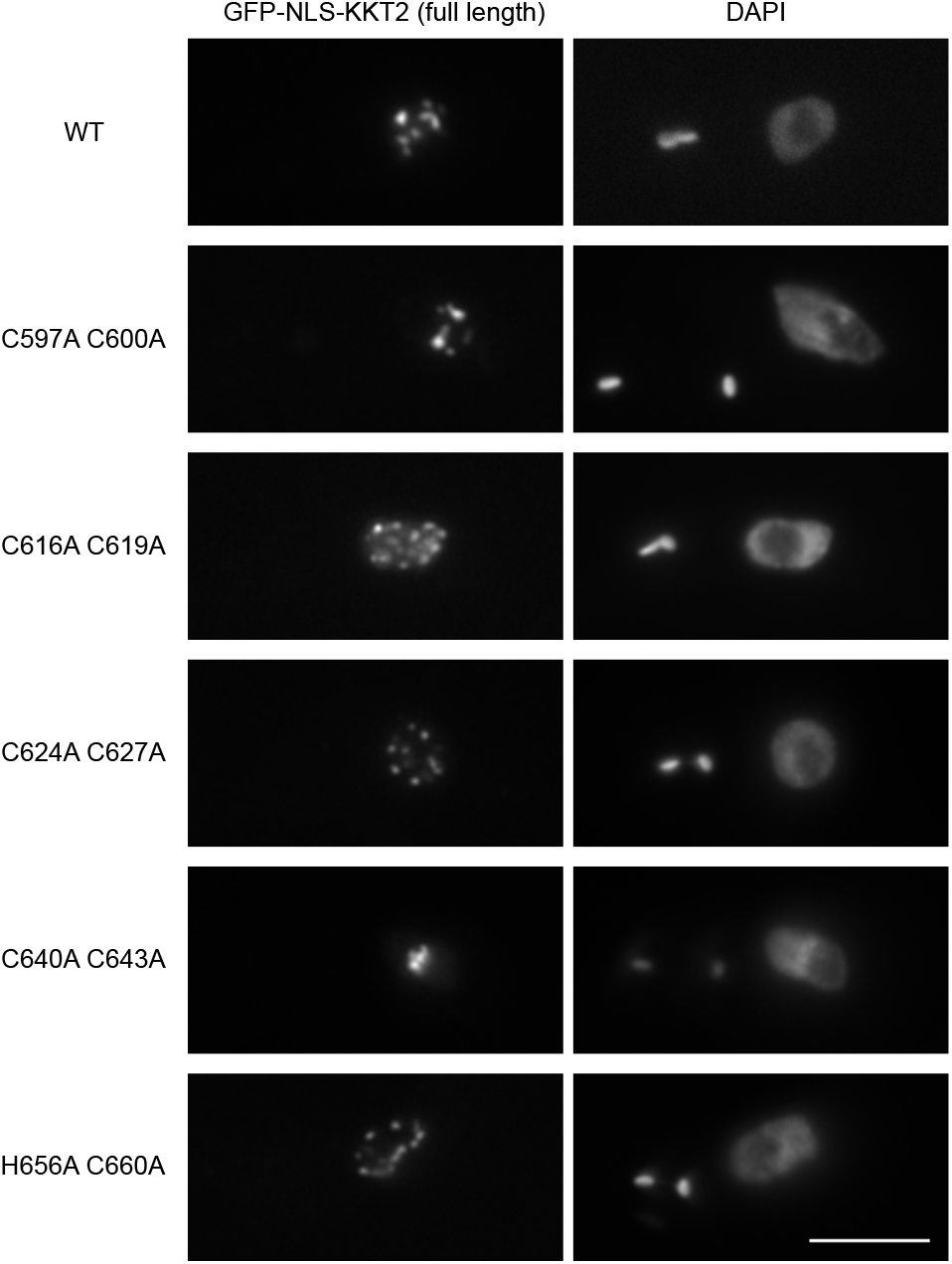
Full-length KKT2 mutants localize at kinetochores. Full-length KKT2 proteins with indicated mutations were ectopically expressed with 10 ng/mL doxycycline. Cell lines, BAP327, BAP365, BAP366, BAP367, BAP368, BAP369. Scale bar, 5 μm.

**Figure S5.**
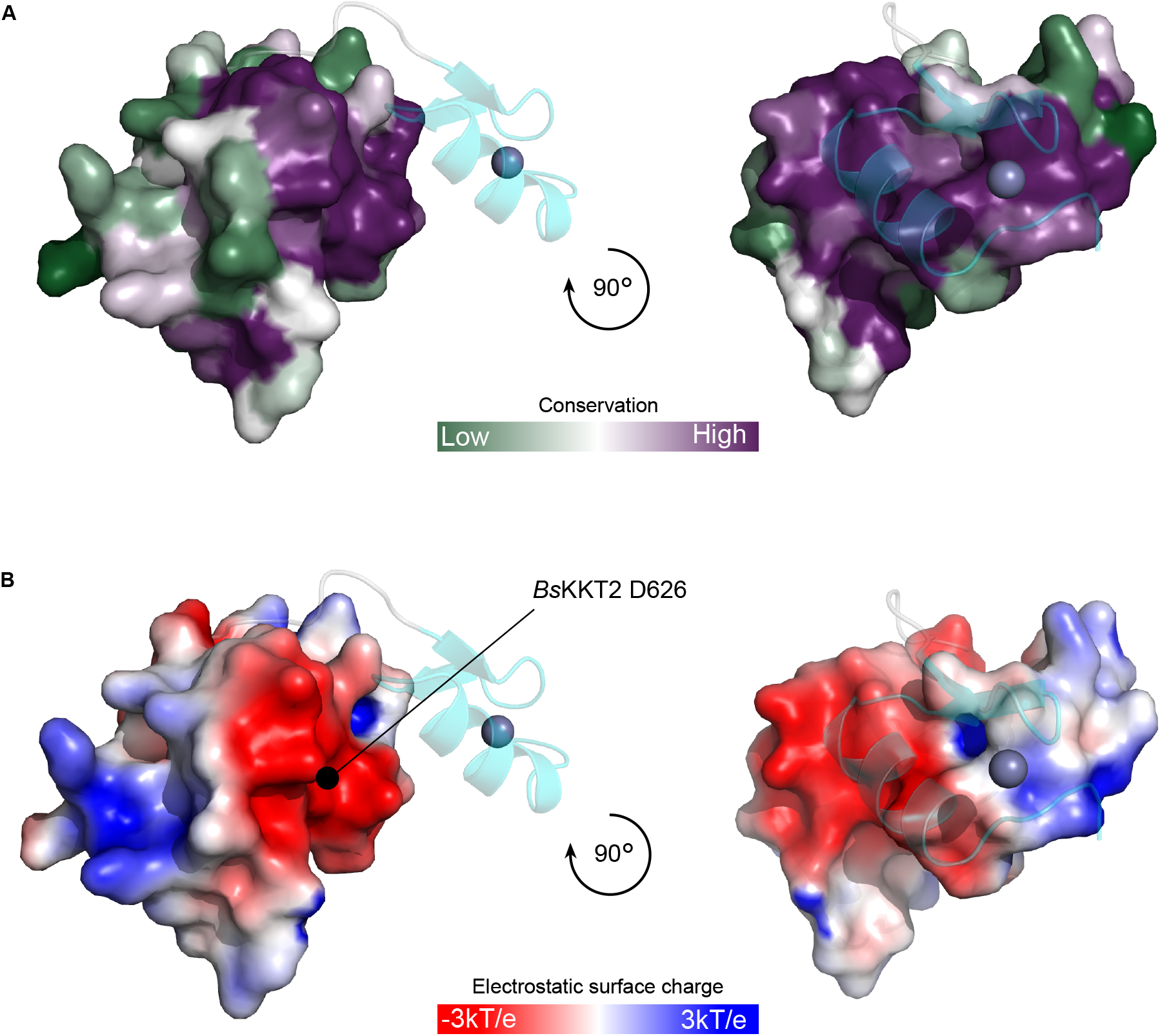
*Bodo saltan*s KKT2 CL domain surface sequence conservation and electrostatic potential reveal the presence of a conserved acidic patch centering on residue D626. (A) Surface sequence conservation of *Bs*KKT2 CL domain using the ConSurf server (Landau et al., 2005; Ashkenazy et al., 2016). The C2H2 zinc finger structure is shown as cartoon representation. (B) Electrostatic surface potential of the *Bs*KKT2 CL domain generated by APBS (Jurrus et al., 2018) reveals the presence of a conserved acidic surface. The location of the residue D626 is marked by a black circle.

**Figure S6.**
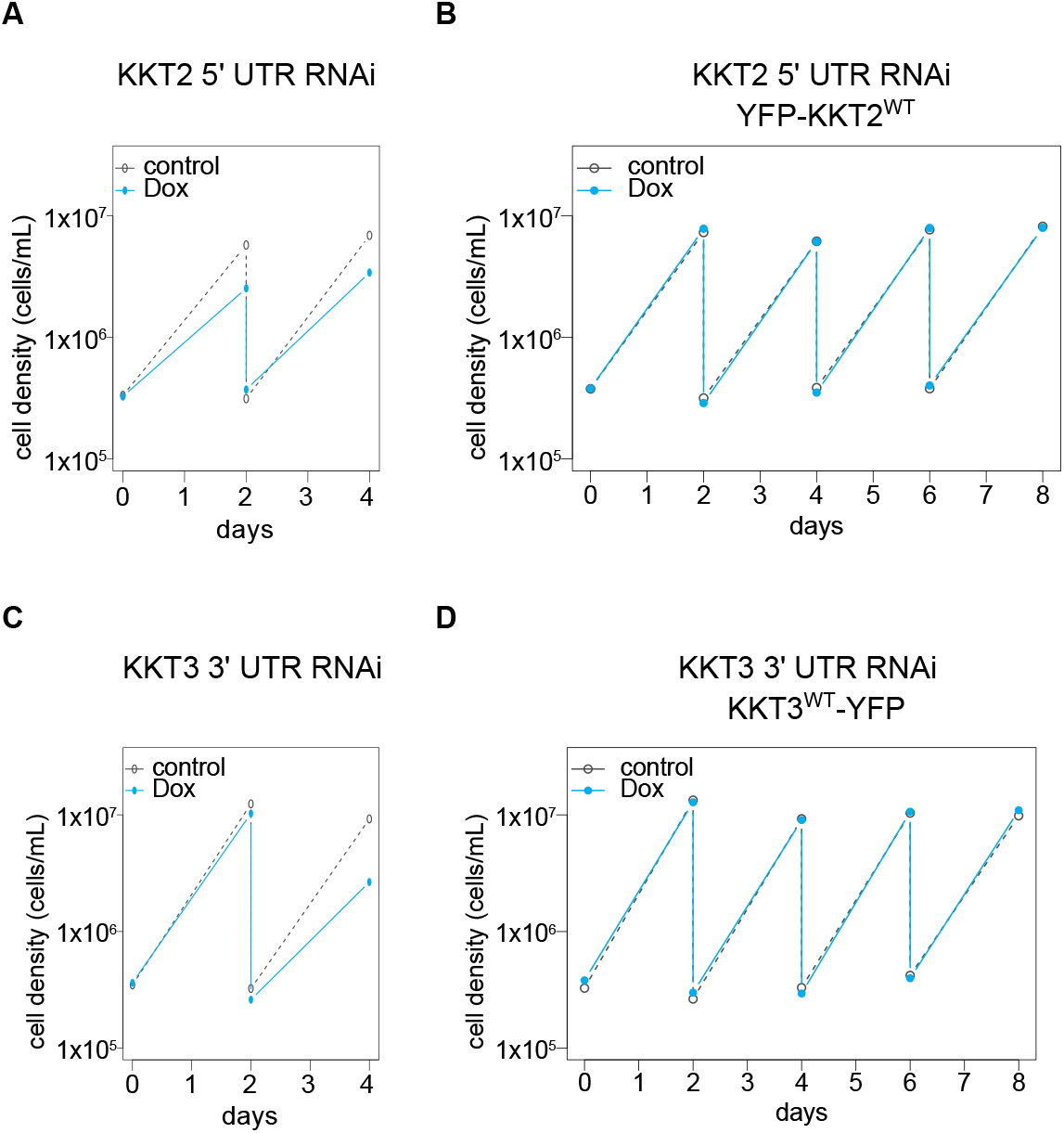
RNAi of KKT2 and KKT3 causes growth defects in *T. brucei*. Growth curves of (A) KKT2 5′UTR RNAi, (B) KKT2 5′UTR RNAi with YFP-KKT2 (resistant to the RNAi), (C) KKT3 3′UTR RNAi, and (D) KKT3 3′UTR RNAi with KKT3-YFP (resistant to the RNAi). 1 μg/mL doxycycline was added to induce RNAi. Controls are uninduced cell cultures. Similar results were obtained from at least two independent experiments. Cell lines, BAP1554, BAP1681, BAP1555, BAP1682.

**Figure S7.**
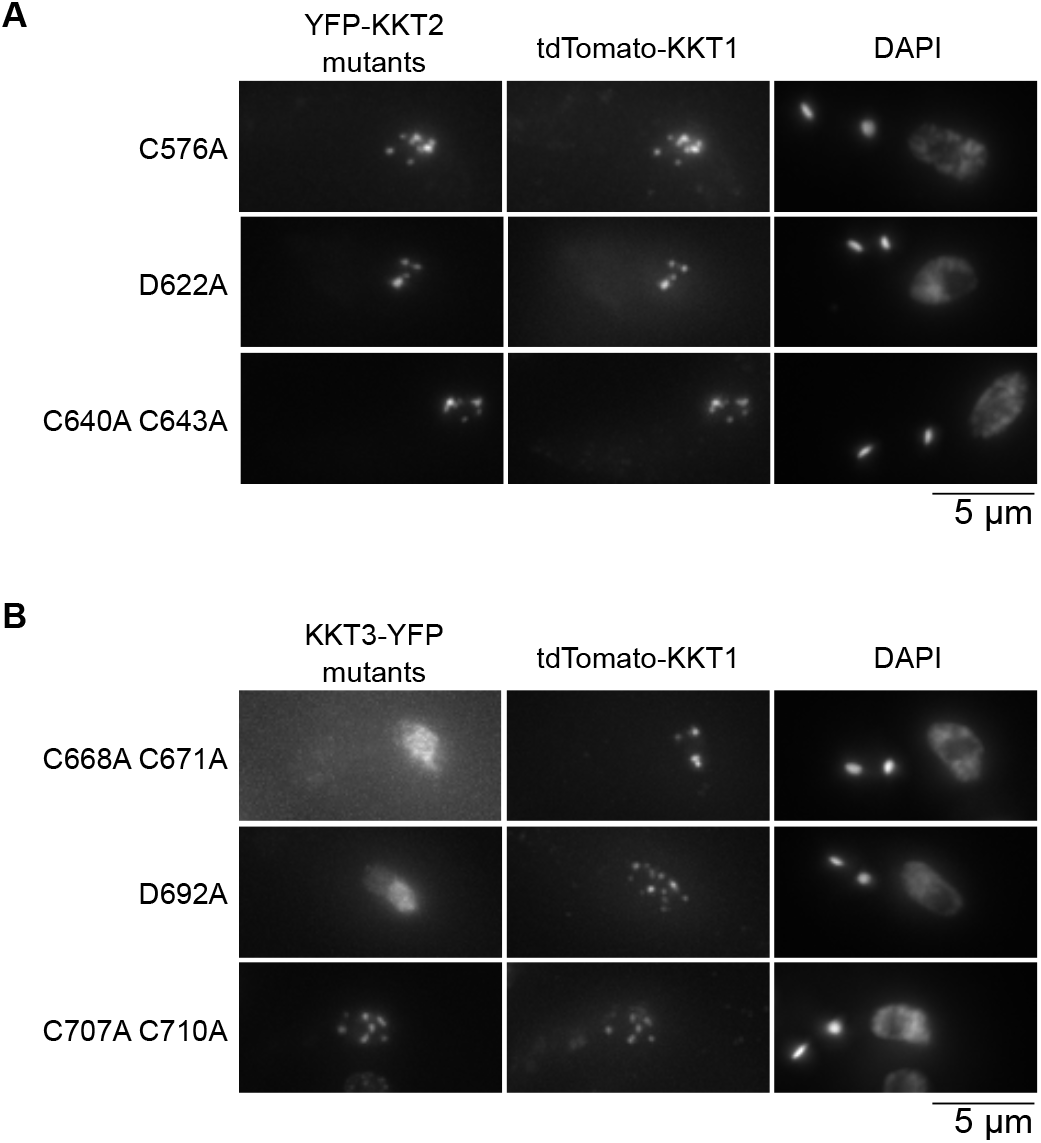
Kinetochore-like dots of KKT2 and KKT3 mutants co-localize with KKT1. (A) Indicated KKT2 mutants expressed at endogenous locus co-localize with a kinetochore marker, tdTomato-KKT1. Cell lines, BAP2036, BAP2033, BAP2035. (B) KKT3^C707A^ ^C710A^ co-localizes with tdTomato-KKT1, while KKT3^C668A^ ^C671A^ and KKT3^D692A^ mutants do not. Cell lines, BAP2037, BAP2038, BAP2034.

